# SMIM1 drives an alternative iron uptake pathway

**DOI:** 10.64898/2026.10.23.761335

**Authors:** Abdelghani Mazouzi, Daniëlle Bianchi, Catrin Lutz, Sarah C. Moser, Guizela Huelsz-Prince, Onno B Bleijerveld, Bim de Klein, Serge Scheffers, Marleen Dekker, Maarten L. Hekkelman, Hendrik A Messal, Alexander Fish, Lodewyk F A Wessels, Jos Jonkers, Thijn R. Brummelkamp

**Affiliations:** Division of Biochemistry, Netherlands Cancer Institute, Amsterdam, Netherlands; Oncode Institute, Utrecht, Netherlands; Division of Molecular Pathology, Netherlands Cancer Institute, Amsterdam, Netherlands; Division of Tumor Biology and Immunology, Netherlands Cancer Institute, Amsterdam, Netherlands; Division of Molecular Carcinogenesis, Netherlands Cancer Institute; Amsterdam, Netherlands; Proteomics Facility, Netherlands Cancer Institute, 1066CX Amsterdam, Netherlands

## Abstract

Iron is essential for numerous biological processes, and its abundance is critical in human cells to prevent anemia or toxic overload^1,2^. Although transferrin receptor-mediated iron uptake is the canonical route^3–5^, human cells also possess an unstudied alternative pathway. To reveal this missing mechanism, we developed a large-scale, gain-of-function approach to assign genes to a catalogue of quantitative cellular traits. Applying this genetic strategy, we identified SMIM1, the antigen responsible for the rare Vel blood group and a transmembrane protein with unknown function, as a key regulator of iron homeostasis. We found that SMIM1 expression completely suppresses the iron deficiency caused by transferrin receptor loss. Mechanistically, SMIM1 mediates iron uptake through clathrin-dependent endocytosis of hemoglobin (Hb). SMIM1 directly binds Hb via its extracellular HKCK motif, leading to its transport to lysosomes and subsequent iron release. In the human population, genetic variants in SMIM1 influence Hb levels and erythrocyte traits. SMIM1-deficient mice display iron deficiency in early erythroid precursors in response to erythropoietic stress, underscoring the physiological relevance of this process.

## Introduction

Iron is crucial for various biological processes, including hemoglobin (Hb) biosynthesis for oxygen transport, mitochondrial function, DNA synthesis, and numerous enzymatic reactions essential for cell survival^2–5^. The human body produces approximately 200 billion red blood cells (RBCs) daily, each containing about 6 g of Hb and 20 mg of iron^4^. To meet this high demand, sophisticated mechanisms have evolved to ensure effective iron acquisition, which primarily involves the transferrin receptor (TFRC)-mediated pathway^2–5^. Defects in iron uptake mechanisms may lead to iron deficiency and anemia. Similarly, iron overload is toxic and equally detrimental^1^. Therefore, iron levels are tightly regulated to maintain homeostasis, and various mechanisms operate to control iron balance^3,4,7^. The primary regulatory network is the Iron-Regulatory Protein (IRP)-Iron-Responsive Element (IRE) system^3^. Cellular iron shortage is sensed through the accumulation of Iron-Regulatory Protein 2 (IRP2), which regulates the post-transcriptional expression of genes involved in iron metabolism, including TFRC^3–5^.

### A novel transferrin receptor-independent pathway for iron uptake

In order to study iron uptake, we inactivated TFRC in HAP1 cells and the liver-derived cell line HepG2. As expected, TFRC loss in HAP1 cells induced iron deficiency, as detected by accumulation of the iron sensor IRP2 (Fig. 1a-b, Extended Data Fig.1a). To our surprise, loss of TFRC did not result in IRP2 upregulation in HepG2 cells (Fig. 1a-c). Importantly, these cells are proficient in sensing iron shortage as iron chelation with deferoxamine (DFO) stabilized IRP2 (Fig. 1c). These observations suggest the presence of an alternative iron acquisition pathway in HepG2 cells that operates independently of the classical transferrin receptor-mediated mechanism and is inactive in HAP1 cells. Interestingly, serum deprivation induces IRP2 stabilization to a similar extent regardless of TFRC status in HepG2 cells. A similar effect is observed following inhibition of endocytosis using Dynasore, a GTPase inhibitor that rapidly inhibits dynamin activity, suggesting that this alternative pathway may mediate iron uptake from serum (Fig. 1d, Extended Data Fig. 1b).

**Figure 1.**
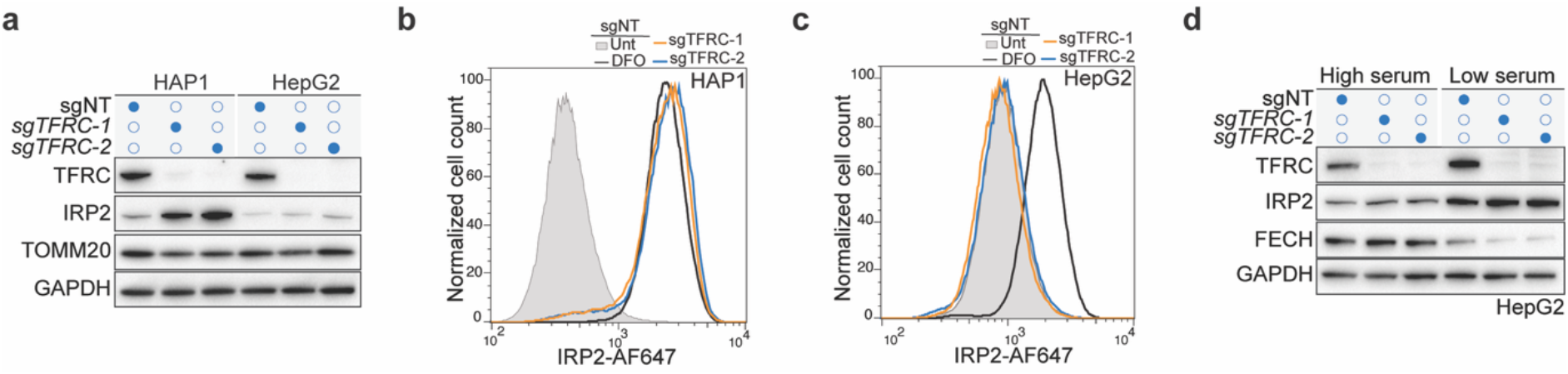
A novel iron uptake pathway independent of the transferrin receptor. (**a-c**) HAP1 or HepG2 wild-type cells were transduced with a non-targeting sgRNA (sgNT) or two independent sgRNAs targeting TFRC. After puromycin selection, whole-cell extracts were subjected to immunoblot analysis using the indicated antibodies, with TOMM20 and GAPDH used as loading controls (a). Filled circles indicate cells expressing the indicated sgRNA, whereas open circles indicate cells lacking the corresponding sgRNA. Alternatively, cells were stained with antibodies against IRP2 and subjected to flow cytometric analysis (b-c). Wild-type cells treated with 20 µM deferoxamine (DFO) for 24 hours were used as a control. (**d**) sgNT or sgTFRC HepG2 cells were cultured in low serum medium (0.5% FCS) or high serum (10% FCS) for 48 hours. Cells were subjected to immunoblot analysis using the indicated antibodies with GAPDH used as a loading control.

### A gain-of-function screening approach to identify the alternative iron uptake pathway

To identify this alternative iron uptake pathway, we applied a genome-wide gain-of-function approach in HAP1 cells. We first developed a gain-of-function strategy that complements the loss-of-function genetic approaches in haploid human cells that we have used for different quantitative cellular phenotypes^8–13^. To this end, we generated a retroviral activating insertional mutagen containing the EF1a promoter, which was chosen after comparing the activity of different promoters in HAP1 cells (Fig. 2a). To address if random integration of this promoter could be used to activate genes in HAP1 cells, wild-type cells were infected and stained using PD-L1 as a marker (Fig. 2b). PD-L1 was chosen as a readout because it is not expressed in HAP1 cells in basal conditions but can be readily induced by interferon-γ (IFN-γ) (Extended Data Fig.1c-d). Subsequently, single clones with high PD-L1 expression were sorted from untreated gene-trapped cells using flow cytometry (Extended Data Fig. 1e). After 10 days of culture, the selected clones displayed a several-hundredfold increase in PD-L1 expression compared to uninfected control cells (Extended Data Fig.1f-g). Next-generation sequencing analysis of retroviral integration sites in the high PD-L1-expressing bulk population revealed an enrichment of sense-orientation insertions in the promoter region of the *CD274* gene, which encodes PD-L1 (Extended Data Fig. 1h), indicating that this vector ensures potent gene activation upon insertion into the promoter region. To test if this approach could be used to robustly link genes to specific cellular phenotypes, we applied this gain-of-function screening approach across a range of cellular processes, including iron metabolism, histone modifications such as H3K27 acetylation and H2A ubiquitination, diverse signaling cascades such as the AKT, mTOR, and p38 pathways, lysosomal biology, tubulin acetylation, poly-ADP-ribosylation, and PD-L1 expression (Fig. 2c, Extended Data Fig. 2a, Table S3). Analysis of insertion enrichment across varying intervals within promoter regions revealed that maximal phenotypic enrichment for most genes occurred in a region spanning −6 kbp upstream of the transcription start site (TSS) up to and including the 5’ UTR. Therefore, this interval was selected as the optimal region to score gene-activating insertions (Extended Data Fig. 2b). The gain-of-function approach allowed the identification of specific regulators (Fig. 2d, Extended Data Fig. 2c, Extended Data Fig. 3a-f) that were not identified using loss-of-function genetics (Fig. 2d, 2g-h, Extended Data Fig. 3a-f), including factors that are redundant (χ^2^ = 189.93, *p*-value = 1.49 × 10^−42^; Fig. 2e) and/or not expressed (Fig. 2f). The screens generated a rich dataset with 4,191 significant outliers (FDR-corrected *p*-value ≤ 0.05 with a Log_2_ mutational index greater than 1 or less than −1) in at least one of the 20 genetic maps (10 for gain-of-function and 10 for loss-of-function phenotypes). We next applied data-driven clustering using a perturbation-perturbation correlation matrix (Extended Data Fig. 4a). For visualization, we employed dimensionality reduction techniques and clustered the data using HDBSCAN (Fig. 2i). Combining both loss- and gain-of-function approaches creates a more comprehensive overview of the factors affecting a defined molecular readout, such as AKT phosphorylation (Fig. 2j, Extended Data Fig. 4b-c). However, despite not including specific phenotypic readouts for mitochondrial processes, vacuolar transport, NuRD complex function or the regulation of RNA metabolic processes, our analysis identified distinct clusters associated with these functions (Fig. 2i). This indicates that our approach can detect cellular changes indirectly caused by disruptions in these pathways, even with a limited set of molecular readouts. The map assigns new regulators to phenotypes such as USP36 acting as modulator of H2A ubiquitination (Extended Data Fig. 5a-e), TMEM253 as a regulator of LAMP1 abundance (Extended Data Fig. 6a-g), and a novel poly-ADP-ribosylation regulatory mechanism operating via the ART1-ENPP1 axis (Extended Data Fig. 7a-k).

**Figure 2.**
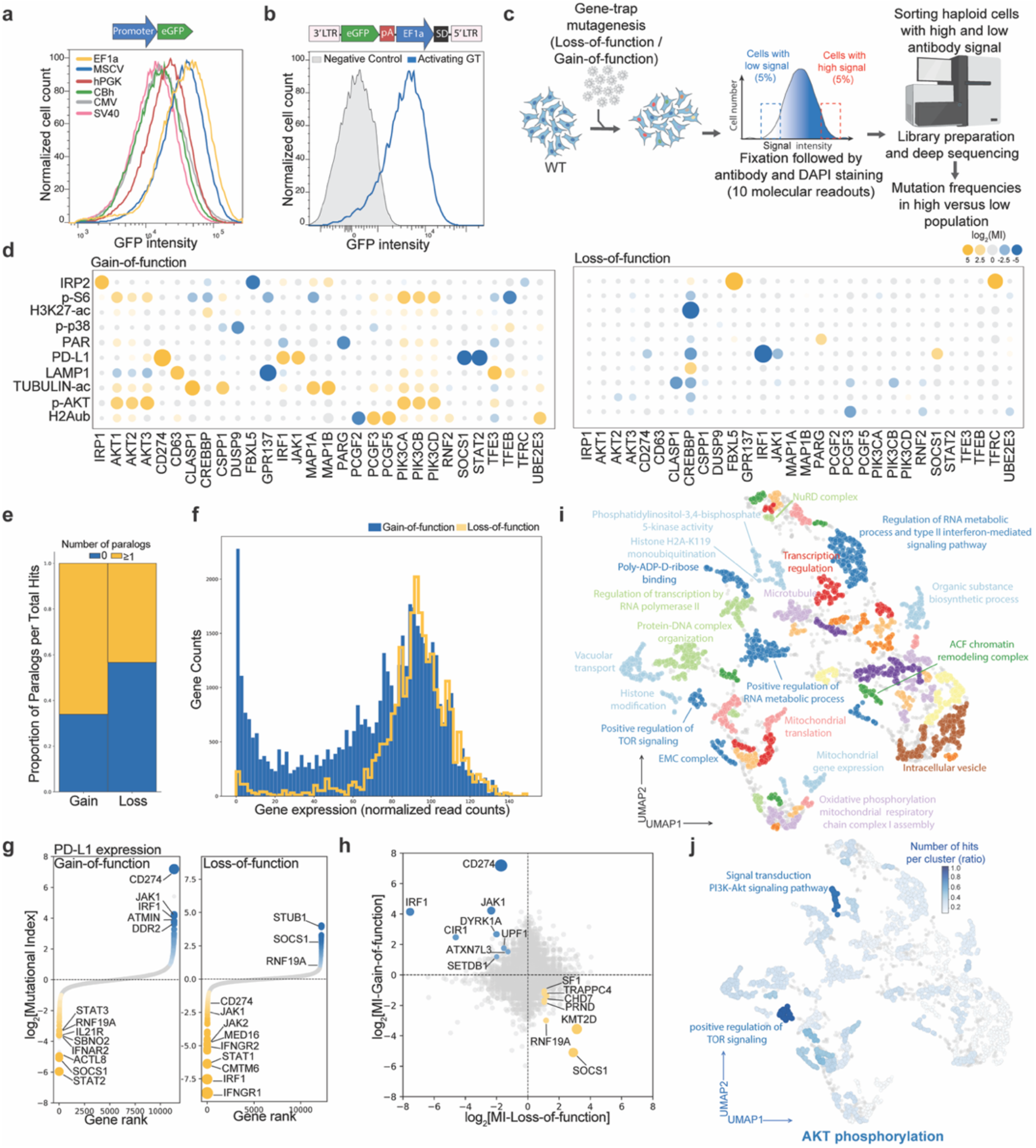
The implementation of gain-of-function screening approach. **(a)** FACS plot displaying the strength of various promoters that were introduced into HAP1 cells by retroviral transduction using GFP as a reporter gene. (**b**) The EF1a promoter was cloned into a retroviral vector to enable activation of genes upon insertion upstream of the ATG start codon and transduced into HAP1 cells. Insertion into active genes leads to GFP expression as an indicator of transduction efficiency. (**c**) Schematic overview of the experimental setup for FACS-based haploid genetic screens using a loss- and a gain-of-function approach and various molecular readouts. (**d**) Bubble plots indicating the effect of selected genes that are known to affect one of the respective phenotypes across the different genetic screens carried out using both gain- and loss-of-function FACS-based screens. Regulators that increase the readout signal upon mutation are colored yellow, while those that decrease the signal are shown in blue. The color contrast is based on the mutation index (MI) (FDR-corrected p ≤ 0.05). Non-significant hits are marked in grey. (**e**) Proportion of number of paralogs identified in the gain- and loss-function screens normalized to the total number of the identified hits. Significance was assessed by a Chi-Square Test (χ2 =189.93, p-value =1.49 × 10^−42^). (**f**) RNA expression levels of the hits that scored in the both gain- and loss-of function screens in HAP1 cells. (**g**) PD-L1 screens conducted using both loss- and gain-of-function haploid genetic under low-dose of IFN-γ treatment (0.5 ng/ml for 24 hours). The data is presented as ranking plots based on mutation index (MI) values. Genes significantly enriched in either channel (two-sided Fisher’s exact test; FDR-corrected p-value ≤ 0.05; total number of unique insertions > 50) are color-coded and top regulators are labeled. (**h**) Scatter plots of the MI value for the loss-of-function PD-L1 screen (x-axis) compared to the gain-of-function (y-axis), with inversely correlated genes labeled. (**i**) UMAP embedding summarizing gene clustering based on aggregated phenotype scores from both loss- and gain-of-function genetic screens using 10 molecular readouts across various biological processes. Colors correspond to manually annotated HDBSCAN clusters containing the labeled gene ontology terms. (**j**) Significant hits from the conducted AKT phosphorylation screens using both loss- and gain-of-function genetics are displayed on the UMAP in (i) with the number of hits scored in each HDBSCAN cluster represented by color intensity.

### SMIM1 drives a transferrin receptor-independent iron uptake pathway

Considering that our gain-of-function approach can activate processes that are normally not active in HAP1 cells, we sought to activate the mechanism of transferrin receptor-independent iron acquisition in HAP1 cells, using IRP2 levels as a sensor for cellular iron shortage (Fig. 3a). We assessed the validity of using IRP2 as a sensor for iron homeostasis by performing both loss-of-function and gain-of-function screens in wild-type HAP1 cells. We identified several known rate-limiting factors for iron metabolism, including TFRC (TFR1), but not its homolog TFR2 (Fig. 3b, Extended Data Fig. 8a), in line with the literature showing that TFR2 has a lower affinity for transferrin and is thought to have a limited role in iron import^14^; instead, it is considered a sensor of transferrin saturation^4,14^. We additionally identified Nuclear Receptor Coactivator 4 (NCOA4), which causes release of stored iron by triggering autolysosomal degradation of ferritin^15^, STEAP3, which reduces iron from its ferric (Fe^3+^) to ferrous (Fe^2+^) form in early endosomes^16^, and the FBXL5-SKP1-CUL1 complex, which regulates IRP2 abundance. These regulators scored in the low channel using the gain-of-function approach and in the high channel of the loss-of-function screens (Fig. 3b, Extended Data Fig. 8a). Furthermore, we identified factors that were identified uniquely using the loss-of-function approach, such as Clathrin Heavy Chain (CLTC) and GTP-binding protein Dynamin 2 (DNM2), which mediate endocytosis of the transferrin-TFRC complex; multiple vacuolar-type ATPases (v-ATPases), which maintain the acidic environment within the early endosome to facilitate the release of iron from transferrin; factors mediating iron trafficking to mitochondria (SLC25A37); and several regulators involved in Fe-S cluster biosynthetic pathways (Extended Data Fig. 8a). These data indicate that genetic screens using IRP2 as readout effectively recapitulate the key factors involved in iron metabolism, and that HAP1 cells primarily rely on the TFRC-mediated iron uptake pathway to acquire iron. To uncover the transferrin receptor-independent iron acquisition pathway, we focused on factors that caused a significant decrease in IRP2 protein levels specifically in TFRC knockout cells but not in wild-type cells using the gain-of-function approach (Fig. 3b-c). This yielded 59 genes (Extended Data Fig. 8b), which were ranked based on their impact on IRP2 levels and the uniqueness of their scores across the 10 molecular readouts used in the gain-of-function screens (Extended Data Fig. 8c-d). The top-scoring gene is Small Integral Membrane Protein 1 (SMIM1), a single-pass membrane protein of 78 amino acids (Fig. 3b-c, Extended Data Fig. 8c-d). SMIM1 is the antigen responsible for the Vel blood group, a rare yet clinically important blood type^17–19^, caused by its extracellular carboxyl terminus that can act as an antigen on RBCs. Importantly, the cellular function of SMIM1 remains unknown.

**Figure 3.**
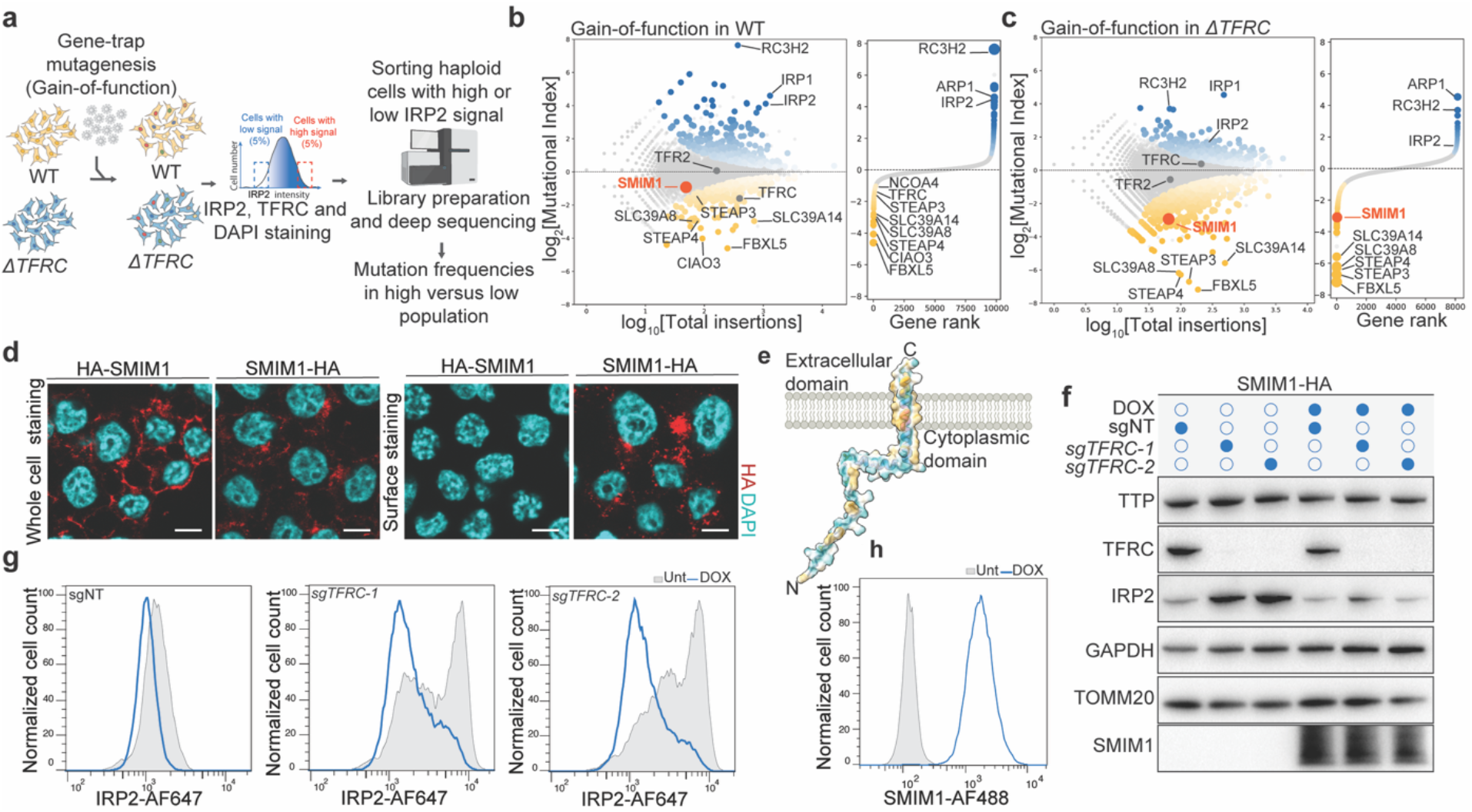
SMIM1 mediates a novel transferrin receptor-independent pathway for iron uptake. (**a**) Schematic overview of the gain-of-function genetic screen in wild-type HAP1 or TFRC-knockout cells using IRP2 as a readout. Briefly, WT or TFRC-knockout cells (sgTFRC-2; *ΔTFRC*) were transduced with the gain-of-function vector as described in the Methods, then surface-stained for TFRC, fixed, and stained for IRP2. TFRC staining was used to exclude non-edited cells from the polyclonal TFRC-knockout population. Haploid TFRC-negative (*ΔTFRC*) or TFRC-positive (WT) cells with high or low IRP2 signals were sorted. Finally, DNA was extracted, and insertions were sequenced and assigned to specific genes. (**b-c**) Fishtail plot of the FACS-based gain-of-function screens using IRP2 as a readout in wild type (b) or TFRC knockout cells (c). The y-axis displays the mutational index (MI) for each gene (see Methods for details). In the fishtail plot, the x-axis indicates the total number of insertions per gene, whereas in the ranking plot, it represents the gene rank. (**d**) Immunofluorescence staining for SMIM1 tagged with an HA tag at either the N or C terminus in HAP1 cells. The cells were surface stained or fixed and permeabilized before staining. Scale bar, 5 μm. (**e**) AlphaFold-predicted structure of SMIM1 with coloring based on hydrophobicity. (**f-h**) HAP1 cells with doxycycline-inducible SMIM1 were transduced with a non-targeting sgRNA (sgNT) or two independent sgRNAs targeting *TFRC*. After puromycin selection and 48 hours of doxycycline induction, whole-cell extracts were subjected to immunoblot analysis using the indicated antibodies, with GAPDH as a loading control (f), or stained with antibodies directed against IRP2 and HA-SMIM1 and subjected to flow cytometric analysis (g-h).

Stable expression of N- or C-terminal HA-tagged SMIM1 in HAP1 cells, analyzed using surface and whole-cell immunofluorescence staining, revealed that the N-terminus of SMIM1 is located in the cytosol, while the C-terminus is exposed outside the plasma membrane (Fig. 3d-e, Extended Data Fig. 8e), consistent with previous reports^20,21^. As suggested by the genetic screens, doxycycline-inducible expression of SMIM1 had minimal effect on IRP2 protein abundance in wild-type cells (Fig. 3f-h, Extended Data Fig. 8f-g). However, the high IRP2 levels observed in TFRC knockout cells were strongly reduced to steady-state levels upon induction of SMIM1, as demonstrated by immunoblot analysis and flow cytometry (Fig. 3f-h, Extended Data Fig. 8h). SMIM1 expression had no effect on TTP (SH3BP4), an important accessory protein involved in the control of TFRC internalization through clathrin-mediated endocytosis^22^, in either wild-type or TFRC-knockout cells (Fig. 3f). These data validate the genetic screen results and indicate that SMIM1 suppresses IRP2 accumulation caused by TFRC loss, suggesting that SMIM1 may mediate an alternative iron acquisition pathway that acts independently of transferrin receptor pathway.

The activity of SMIM1 relies on clathrin-mediated endocytosis, as dynasore completely abolished the action of SMIM1 in TFRC-knockout cells (Fig. 4a-b). A similar effect was observed with bafilomycin A1 (BafA1), a potent macrolide that inhibits the v-ATPase pump and disrupts lysosomal acidification (Fig. 4b). Iron chelation by DFO also abolished the effects of SMIM1 expression in TFRC knockout cells, demonstrating that the phenotype is not caused by interference with iron sensing (Fig. 4b). Consistent with the importance of clathrin-mediated endocytosis in this process, SMIM1 was readily internalized (Extended Data Fig. 9a) and showed colocalization with clathrin (Fig. 4c). These experiments suggest that SMIM1 facilitates TFRC-independent iron uptake via clathrin-mediated endocytosis and lysosomal acidification, indicating a potential role for SMIM1 in the uptake of iron or iron-carrying molecules from the external environment.

**Figure 4.**
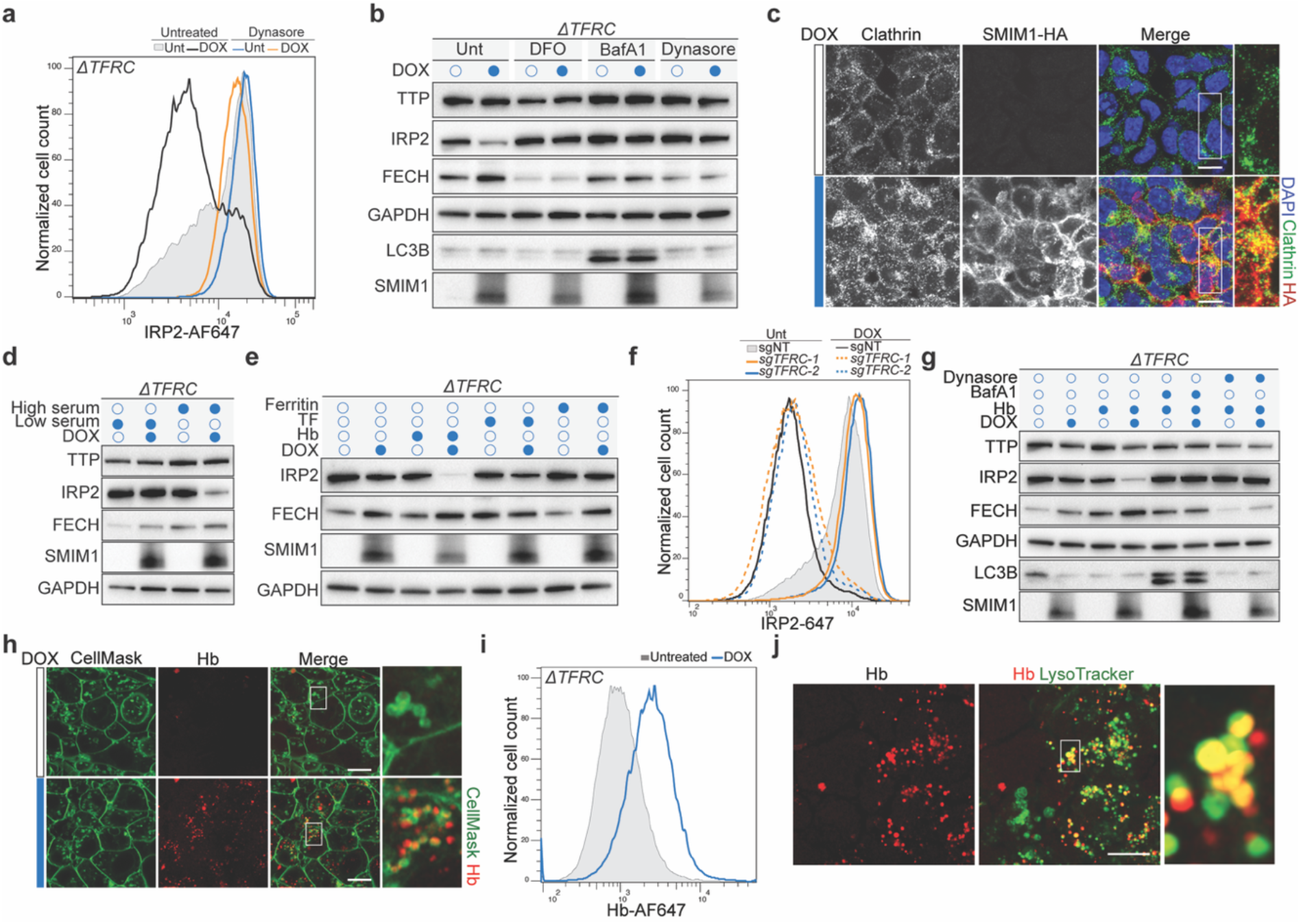
SMIM1 mediates the uptake of Hb. (**a**) TFRC knockout HAP1 cells were treated with doxycycline or left untreated for 24 hours to induce SMIM1 expression, followed by 10 µM dynasore treatment for an additional 24 hours or untreated. The cells were immunostained using antibodies directed against IRP2 and subjected to flow cytometry analysis. (**b**) TFRC-knockout HAP1 cells were treated with doxycycline or left untreated for 24 hours to induce SMIM1 expression, followed by treatment with 10 µM deferoxamine (DFO), 10 nM Bafilomycin A1 (BafA1), or 10 µM dynasore for an additional 24 hours or left untreated. Whole-cell extracts were subjected to immunoblot analysis using the indicated antibodies, with GAPDH as a loading control. (**c**) TFRC-knockout HAP1 cells were treated with doxycycline for 24 hours to induce SMIM1 expression or left untreated as a control, followed by immunofluorescence staining for C-terminal HA-tagged SMIM1 and clathrin heavy chain (CLTC). DNA was stained with DAPI. The open box indicates untreated cells, whereas the filled box indicates cells treated with doxycycline (DOX), as described above. Scale bar, 10 μm. (**d**) TFRC-knockout HAP1 cells were treated with doxycycline to induce SMIM1 expression or left untreated as a control under either low-serum (0.5% FCS) or high-serum (10% FCS) conditions for 48 hours. Whole-cell extracts were analyzed by immunoblot using the indicated antibodies, with GAPDH as a loading control. (**e**) SMIM1 was induced in TFRC knockout HAP1 cells using doxycycline treatment for 16 hours or left untreated in low serum (0.5% FCS). Subsequently, the cells were cultured with 10 µg/ml Hb, 10 µg/ml transferrin (TF), or 10 ng/ml ferritin for an additional 24 hours. Immunoblot analysis was then performed on whole-cell extracts using the indicated antibodies, with GAPDH as the loading control. (**f**) Wild type HAP1 cells carrying doxycycline-inducible SMIM1 were transduced with either a non-targeting sgRNA (sgNT) or two independent sgRNAs targeting TFRC. SMIM1 expression was induced by treatment with doxycycline for 24 hours or left uninduced as a control in low serum. Cells were subsequently cultured in low serum medium supplemented with 10 µg/ml hemoglobin (Hb) for an additional 24 hours. Cells were then fixed, immunostained for IRP2, and analyzed by flow cytometry. (**g**) TFRC-knockout HAP1 cells were exposed to doxycycline for 16 hours to induce SMIM1 expression or left untreated, then cultured in the presence or absence of 10 µg/ml Hb, 10 nM Bafilomycin A1 (BafA1), or 10 µM dynasore for an additional 24 hours in low serum. Whole-cell extracts were subjected to immunoblot analysis using the indicated antibodies, with GAPDH as a loading control. (**h**) TFRC-knockout HAP1 cells were exposed to doxycycline for 24 hours to induce SMIM1 expression or left untreated, then cultured in the presence of fluorescent Hb (Hb-AF647, 1 µg/ml) for 6 hours. The cells were subsequently analyzed by live-imaging confocal microscopy to detect Hb-AF647 and cell membranes stained with CellMask. Scale bar, 10 μm. (**i**) Flow cytometry analysis to detect Hb-AF647 was performed under the same experimental conditions described in (h). (**j**) Representative image of the co-localization of Hb-AF647 with the lysosome marker (LysoTracker) in TFRC-knockout HAP1 cells. Scale bar, 10 μm.

### SMIM1 modulates iron metabolism through hemoglobin uptake

To study if the activity of SMIM1 would require serum components, we cultured TFRC knockout cells in either low or high serum media, with or without SMIM1 expression. As expected, SMIM1 expression led to a significant reduction in IRP2 levels in cells cultured in high serum, whereas this effect was suppressed in cells cultured in low serum (Fig. 4d and Extended Data Fig. 9b). This effect is unlikely to be through the direct transport of free iron or heme, as supplementation of the low serum media with iron in the form of ferric ammonium citrate (FAC) or hemin chloride had a similar effect on IRP2 levels, regardless of SMIM1 status (Extended Data Fig. 9c-d). This finding supports the idea that SMIM1-dependent uptake involves iron-carrying proteins, in line with its requirement for clathrin-mediated endocytosis and lysosomal acidity. Therefore, we analyzed the composition of fetal bovine serum used as a supplement to our medium using mass spectrometry. We first performed ammonium sulfate precipitation to remove the highly abundant bovine serum albumin (Extended Data Fig. 9e), which could hinder the identification of low-abundant serum constituents. The supernatant of the 40% ammonium sulfate precipitated serum retained the effect on IRP2 levels observed upon SMIM1 expression (Extended Data Fig. 9f), and mass spectrometric analysis revealed several iron-carrying proteins, including transferrin (TF) and different subunits of Hb, such as HBA, HBB, and fetal Hb (fHb) (Extended Data Fig. 9g). Based on these results, we decided to test the effect of human Hb, transferrin (TF) and ferritin, a protein that sequesters iron and is secreted extracellularly^5^. Supplementation with transferrin or ferritin did not affect IRP2 levels of TFRC knockout cells grown in low serum in the presence or absence of SMIM1 (Fig. 4e). In contrast, SMIM1-expressing cells supplemented with human Hb exhibited a striking decrease in IRP2 levels and stabilization of FECH, the final enzyme of the heme biosynthesis pathway, upon SMIM1 expression (Fig. 4e, g). The effect of SMIM1 expression on IRP2 stability was also confirmed using flow cytometry (Extended Data Fig. 9h). A similar effect was observed with bovine Hb (Extended Data Fig. 9i). This effect on IRP2 is independent of TFRC, as culturing both wild type and TFRC knockout cells in low serum medium supplemented with human Hb resulted in a significant reduction in IRP2 levels upon SMIM1 expression (Fig. 4f). Treatment with dynasore suppresses the effect of SMIM1 on IRP2, indicating that SMIM1 mediates the uptake of Hb through clathrin-mediated endocytosis (Fig. 4g). Moreover, lysosomal acidity is required for the effect of SMIM1 on IRP2 in the presence of Hb (Fig. 4g), as we previously showed in the serum experiment (Fig. 4b). In line with this, loss-of-function screens using IRP2 as a readout in doxycycline-inducible SMIM1 HAP1 cells cultured in low serum in the presence of Hb and in the presence or absence of doxycycline, highlighted SMIM1-specific factors related to lysosomal lumen acidification, trafficking, and Fe-S cluster assembly, confirming our previous findings (Extended Data Fig. 10a-d).

To study SMIM1-mediated Hb uptake, doxycycline-inducible SMIM1 TFRC-knockout HAP1 cells were cultured with or without doxycycline, and uptake of fluorescently labeled Hb was analyzed by microscopy and flow cytometry. SMIM1 expression led to a strong uptake of labeled Hb (Fig. 4h-i), which was internalized and colocalized with lysosomes (Fig. 4j). Upon endocytosis inhibition by dynasore, Hb remains at the cytoplasmic membrane, where SMIM1 is localized, in contrast to the untreated control cells (Extended Data Fig. 9k-l).

Hb is released into the plasma during intravascular hemolysis, which accounts for over 10% of daily RBC turnover. Free Hb is then captured by the acute-phase protein haptoglobin^23^, leading to the formation of a haptoglobin-Hb complex, which is subsequently removed from the circulation by the scavenger receptor CD163 expressed on macrophages^24^. Although CD163 is not expressed in HAP1 cells (Extended Data Fig. 9m), we wanted to determine whether SMIM1 operates through a haptoglobin-dependent pathway. We therefore cultured doxycycline-inducible SMIM1 TFRC-knockout HAP1 cells with or without doxycycline, in the presence of Hb with or without haptoglobin. Interestingly, the addition of haptoglobin mitigated the reduction in IRP2 observed upon SMIM1 expression (Extended Data Fig. 9n) and completely inhibited Hb uptake (Extended Data Fig. 9o), demonstrating that haptoglobin binding to Hb competes with its internalization through SMIM1. Thus, SMIM1 facilitates Hb uptake through clathrin-mediated endocytosis via a mechanism independent of the canonical haptoglobin-CD163 pathway, leading to subsequent iron release.

### SMIM1 directly binds hemoglobin via its conserved extracellular HKCK motif

To address if SMIM1 directly interacts with Hb, we first incubated HAP1 cells loaded with the extracellular domain of SMIM1 (SMIM1-ECD-WT, corresponding to the C-terminal 15 residues of SMIM1) with labeled Hb or control and analyzed them by flow cytometry. Cells with SMIM1-ECD-WT showed strong staining with Hb compared to the control (Fig. 5a), in a dose-dependent manner (Extended Data Fig. 11a). This effect depended on the last four conserved amino acids of SMIM1, as their mutation to alanine (HKCK to AAAA; SMIM1-ECD-4A) completely abolished staining with labeled Hb (Fig. 5a, Extended Data Fig. 8e). Immunofluorescence analysis showed that the extracellular domain of SMIM1 is sufficient to decorate cell membranes with labeled Hb (Fig. 5b, Extended Data Fig. 11d). This phenotype was observed across various cell types (Fig. 5c, Extended Data Fig. 11b-c). Consistent with this, *in vitro* reconstituted liposomes loaded with SMIM1-ECD-WT showed strong staining with labeled Hb compared to the control, an effect that was completely suppressed in liposomes with the SMIM1-ECD-4A mutant (Fig. 5d). We further confirmed the direct interaction between the extracellular domain of SMIM1 and Hb by performing a surface plasmon resonance (SPR) experiment. Hb showed specific binding to the immobilized extracellular domain of wild-type SMIM1, whereas transferrin exhibited no binding (Fig. 5e-f). This interaction depended again on the last four conserved amino acids, as Hb showed no binding to SMIM1-ECD-4A (Fig. 5e-f). Interestingly, SMIM1 also binds to and mediates uptake of different Hb variants, including sickle cell Hb (HbS), but not myoglobin (Mb), highlighting its specificity for Hb (Fig. 5g-h, Extended Data Fig. 11e-l). Additionally, isothermal titration calorimetry showed that a single Hb molecule binds to 16 molecules of the extracellular domain of SMIM1 (SMIM1-ECD-WT), suggesting that SMIM1 may require multimerization for efficient Hb binding (Extended Data Fig. 12a-b). Overall, these data indicate that SMIM1 mediates Hb uptake through a direct physical interaction via its conserved HKCK motif and requires multimerization for efficient Hb binding.

**Figure 5.**
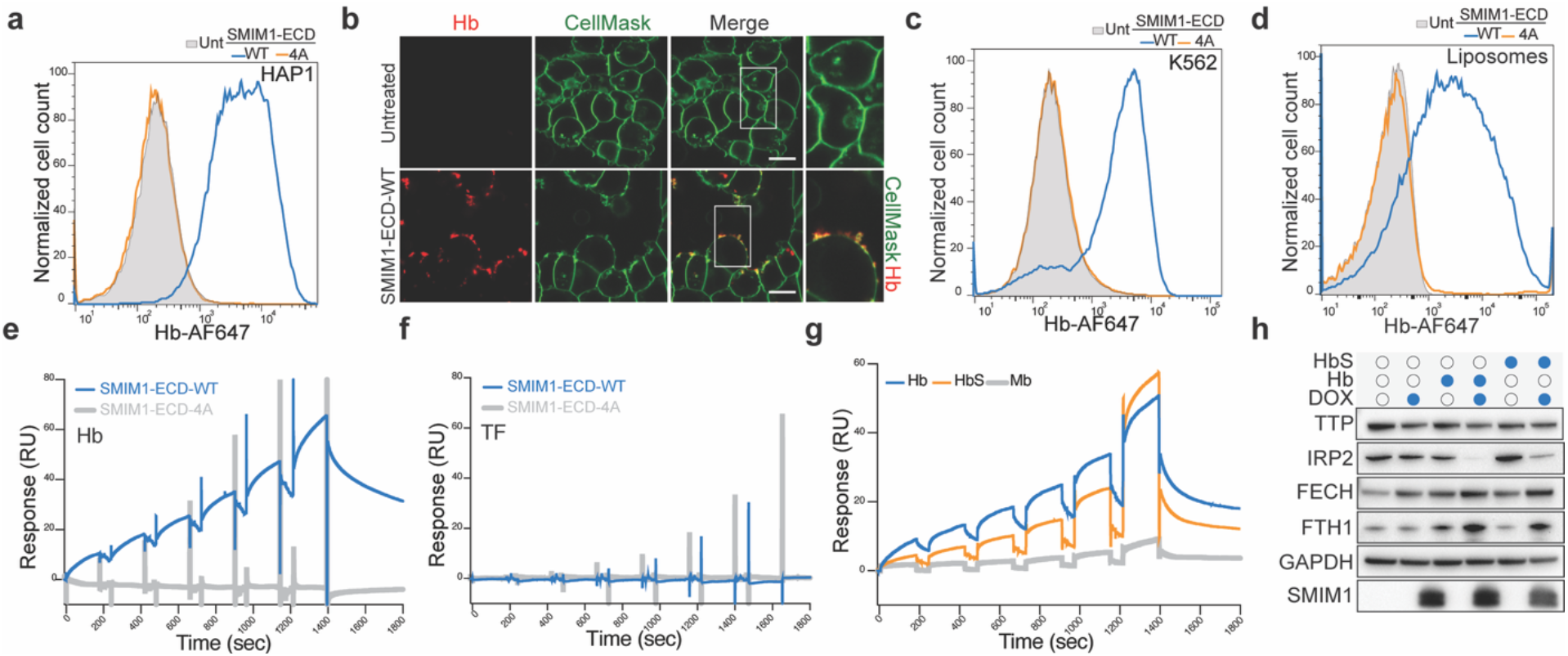
Binding of SMIM1 to Hb. **(a)**Wild-type HAP1 cells were incubated for 2 hours with 1 µg/ml labeled Hb (Hb-AF647) in the presence of either DMSO, 12.5 μM wild-type SMIM1 extracellular domain (SMIM1-ECD-WT; C-terminal 15 amino acids of SMIM1), or the alanine-mutated HKCK motif variant (SMIM1-ECD-4A). The cells were then washed, harvested, and subjected to flow cytometry analysis for Hb fluorescence. **(b)** Confocal microscopy to detect fluorescent Hb (Hb-AF647, 1 µg/ml) and CellMask to stain the plasma membrane in wild type HAP1 cells incubated with either DMSO or 12.5 μM SMIM1-ECD-WT for 2 hours in the presence of 1 µg/ml Hb-AF647. Scale bar, 10 μm. (**c**) A similar experimental setup as in (a) was performed in K562 cells. (**d**) *In vitro* reconstituted liposomes were incubated for 2 hours at 37°C with either DMSO or 12.5 μM SMIM1-ECD-WT or SMIM1-ECD-4A, in the presence of labeled hemoglobin (Hb-AF647). The liposomes were then washed and subjected to flow cytometry to measure Hb fluorescence. (**e-f**) Surface plasmon resonance (SPR) analysis of binding of human Hb (e) or transferrin (TF) as a control (f) to the immobilized wild-type or alanine-mutant extracellular domain of SMIM1. (**g**) Binding of human Hb, sickle cell Hb (HbS), or myoglobin (Mb) to the immobilized wild type of the extracellular domain of SMIM1, measured by SPR. (**h**) SMIM1 expression was induced in TFRC knockout HAP1 cells by treatment with doxycycline for 16 hours or left uninduced as a control. Cells were subsequently cultured in low-serum medium (0.5% FCS) supplemented with either 10 µg/ml hemoglobin (Hb) or 10 µg/ml sickle hemoglobin (HbS) for an additional 24 hours. Whole cell extracts from the conditions were then analyzed by immunoblot using the indicated antibodies, with GAPDH as a loading control.

### Physiological role of SMIM1-dependent iron uptake

SMIM1 is reported to be highly expressed in RBC precursors and erythroblasts^17^. To establish the biological relevance of SMIM1-mediated Hb uptake *in vivo*, we analyzed the maturation stages of differentiating erythroblasts in the bone marrow of wild-type (*Smim1*^*+/+*^) and SMIM1 knockout (*Smim1*^*−/−*^) mice. These mice were left unchallenged or subjected to chemically induced hemolytic anemia by subcutaneous injection of the oxidant phenylhydrazine (PHZ), and bone marrow composition was analyzed 72 hours later. (Fig. 6a, Extended Data Fig. 13a-b). Under unchallenged conditions, no differences were observed between *Smim1*^*+/+*^ and *Smim1*^*−/−*^ mice in the distribution of erythroid differentiation stages, including proerythroblasts (ProE), basophilic (BasoE), polychromatophilic (PolyE), and orthochromatophilic (OrthoE) erythroblasts (Fig. 6b-c, Extended Data Fig. 13c). In response to PHZ-induced erythropoietic stress and hemolytic anemia, both *Smim1*^*+/+*^ and *Smim1*^*−/−*^ mice showed a strong enrichment of early erythroid precursors (ProE and BasoE) relative to PolyE and OrthoE in the bone marrow (Fig. 6d-g, Extended Data Fig. 13d-e). Interestingly, under PHZ-treated conditions, *Smim1*^*−/−*^ mice accumulated erythroid precursors in the bone marrow with higher CD71 (TFRC) expression than wild-type controls (Fig. 6h). This population of cells closely resembled BasoE (Extended Data Fig. 14a-d) but were distinguished by higher levels of CD71, which might be a consequence of iron shortage as immunoblot analysis showed that these cells had high IRP2 levels (Fig. 6i, Extended Data Fig. 14b). This suggests that BasoE cells may experience iron deficiency in the absence of SMIM1-mediated Hb uptake. Indeed, BasoE cells are capable of taking up fluorescently labeled Hb, and this uptake was reduced in SMIM1-deficient mice (Fig. 6j-k). Together, these data suggest that BasoE cells display SMIM1-mediated Hb uptake and that under conditions of erythropoietic stress, these cells use both SMIM1 and TFRC to support enhanced erythropoiesis and prevent iron deficiency.

**Figure 6.**
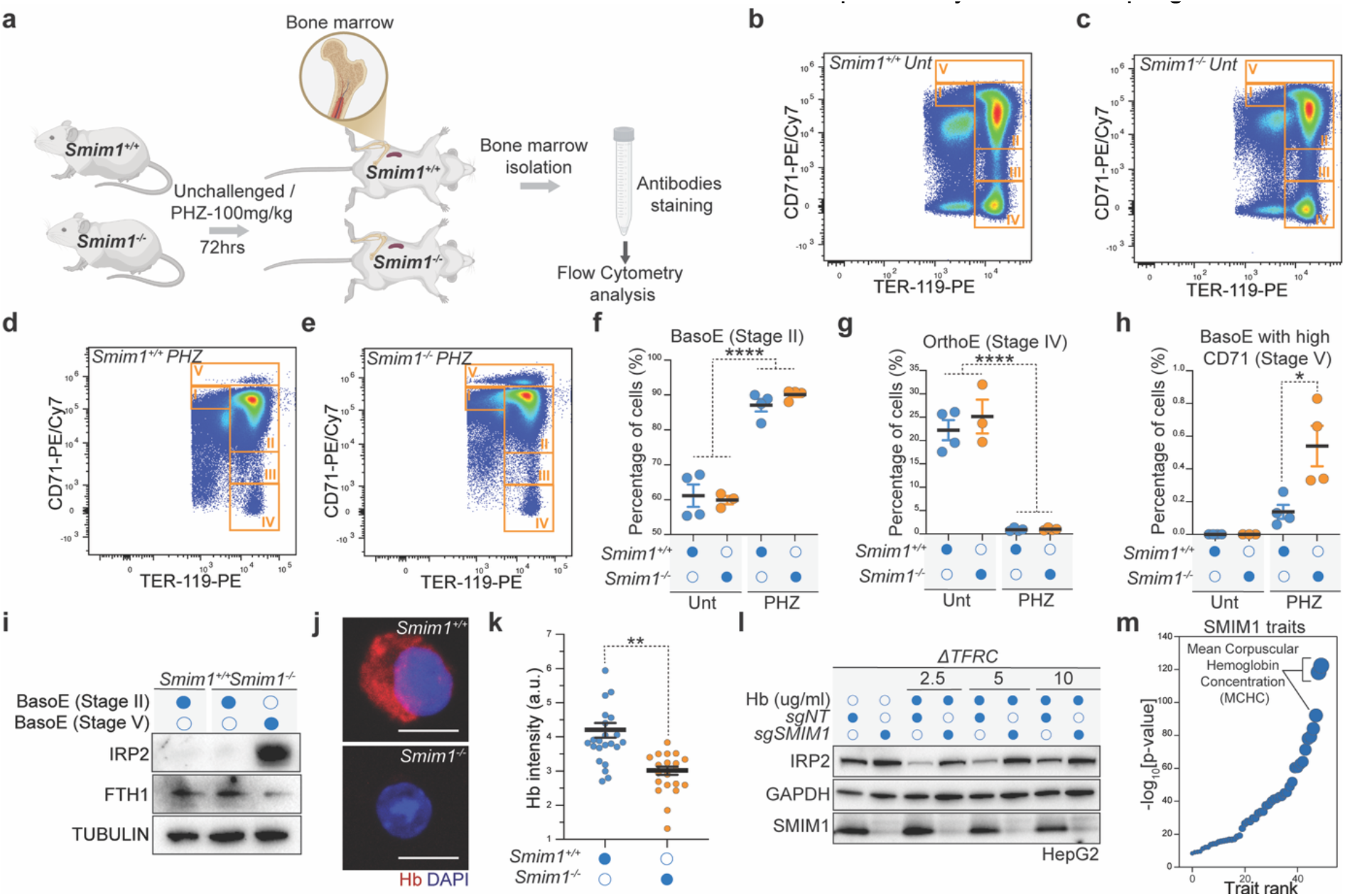
Physiological relevance of SMIM1-dependent iron uptake. **(a)** Schematic overview of the experimental setup for the *in vivo* mouse experiment. Wild type control (*Smim1*^*+/+*^) and SMIM1 knockout (*Smim1*^*−/−*^) mice were left unchallenged or were subjected to a single subcutaneous injection of phenylhydrazine (PHZ, 100 mg/kg). Bone marrow was isolated, and its composition was analyzed by flow cytometry 72 hours later. (**b-e**) Flow cytometry density plots of the bone marrow erythropoietic populations from representative mice: *Smim1*^*+/+*^ (b) and *Smim1*^*−/−*^ (c) under unchallenged conditions, and *Smim1*^*+/+*^ (d) and *Smim1*^*−/−*^ (e) after PHZ treatment. (**f-h**) Frequencies (%) of erythroid precursors, BasoE (f), OrthoE (g), and BasoE with high CD71 (h), in the bone marrow of *Smim1*^*+/+*^ and *Smim1*^*−/−*^ mice under unchallenged conditions (*n = 4*) or after PHZ-induced anemia (*n = 4*). Results are shown as mean ± SEM and were analyzed by a two-sided t-test. *P < 0.05, ****P < 0.0001. (**i**) *Smim1*^*+/+*^and *Smim1*^*−/−*^ mice were challenged with PHZ, and BasoE stage II and stage V cells were isolated and analyzed by immunoblot using the indicated antibodies, with tubulin as a loading control. (**j**) Isolated BasoE from *Smim1*^*+/+*^ and *Smim1*^*−/−*^ mice under unchallenged conditions were cultured in the presence of labeled Hb for 4 hours. The cells were then fixed and analyzed by confocal microscopy to detect fluorescent Hb (Hb-AF647). DNA was stained with DAPI. Scale bar, 5 μm. (**k**) Quantification of Hb-AF647 intensities in more than 20 isolated BasoE per condition. Values represent mean intensity per cell, and the *p*-value was calculated using a two-sided t-test. **P < 0.01. (**l**) TFRC knockout HepG2 cells were transduced with either a non-targeting sgRNA (sgNT) or sgRNAs targeting SMIM1 and cultured in low serum medium with different concentrations of Hb for 48 hours. Whole-cell extracts were subjected to immunoblot analysis using the indicated antibodies, with GAPDH serving as a loading control. (**m**) Ranking plot depicting the association of SMIM1 with blood cell traits based on GWAS studies. The 3 highest-ranking associations relate to mean corpuscular hemoglobin concentration (MCHC), the concentration of hemoglobin relative to the volume occupied by red blood cells (P = 4.09 × 10^−96^). Summary statistics were downloaded from the NHGRI-EBI GWAS Catalog^6^ on 08/08/2024.

Our initial observations suggested the existence of an alternative iron acquisition pathway in liver-derived HepG2 cells, which functions independently of the classical transferrin receptor-mediated mechanism (Fig. 1a-d). Because SMIM1 is expressed in HepG2 cells, we genetically inactivated SMIM1 in TFRC knockout HepG2 cells. Interestingly, SMIM1-depleted cells displayed elevated IRP2 levels compared to control cells in the presence of Hb (Fig. 6l, Extended Data Fig. 11m). These findings suggest that the SMIM1-mediated alternative pathway for iron uptake is potentially active in the liver, which might aid in the clearance of free Hb from the circulation and ensure iron recycling.

## Discussion

Here, we identified an uncharacterized iron uptake pathway that acts independently of the classical transferrin receptor route. This process is mediated by SMIM1, a small protein underlying the Vel blood group antigen, which binds Hb via its extracellular HKCK motif and enables its uptake through clathrin-mediated endocytosis, leading to lysosomal Hb degradation and subsequent iron release. The importance of this alternative pathway is twofold. First, this pathway may serve as an iron recycling mechanism, as Hb represents a major iron reservoir and most circulating iron derives from RBC turnover, especially in response to hematopoietic stress^1,7,25^. The SMIM1 pathway may play a crucial role in this context, as SMIM1 loss in mice leads to iron deficiency in early erythroid precursors in response to erythropoietic stress. Second, Hb is a potentially tissue-damaging molecule due to its highly reactive heme groups and is released into the plasma through intravascular hemolysis^26^. This process can be accelerated in the context of infectious diseases such as malaria or inherited disorders such as sickle cell anemia, leading to serious pathological complications. The SMIM1 pathway may therefore contribute to Hb clearance and prevent its toxicity under conditions of increased circulating Hb. Indeed, we found that SMIM1 binds to and mediates the uptake of sickle cell hemoglobin (HbS).

The canonical pathway for Hb clearance from the circulation involves the haptoglobin-CD163 pathway^24^. However, haptoglobin- and CD163-knockout mice are viable, and the clearance of free plasma Hb is not compromised compared with wild-type mice^27,28^, suggesting the existence of alternative Hb uptake mechanisms. Earlier studies in rat hepatocytes support this possibility^29^. Interestingly, we found that the SMIM1 pathway works independently of the haptoglobin-CD163 pathway and that inactivation of the endogenous SMIM1 gene in a hepatocyte-derived cell line induces iron deficiency. These findings suggest that SMIM1 may represent one such alternative Hb clearance pathway, although further *in vivo* studies are required.

The relationship between SMIM1 and the haptoglobin-CD163 system may depend strongly on physiological context. Under normal conditions, circulating free Hb is expected to be limited by the high scavenging capacity of the haptoglobin-CD163 system^23,24,27^. Indeed, our finding that haptoglobin prevents SMIM1-mediated Hb uptake suggests that SMIM1 is unlikely to compete efficiently with haptoglobin for Hb when circulating haptoglobin is abundant. During accelerated intravascular hemolysis, such as in sickle cell anemia or malaria infection, however, Hb/heme scavenging mechanisms can become saturated, resulting in the accumulation of extracellular Hb. Under these conditions, SMIM1 may provide an additional route for Hb clearance and iron recovery. In line with this, SMIM1 deficiency causes an iron-deficiency phenotype in early erythroid precursors in response to chemically induced hemolytic anemia. This highlights the importance of the SMIM1 pathway when iron demand is increased and extracellular Hb becomes available.

Although SMIM1 loss in mice under steady-state conditions did not result in a significant phenotype, analysis of genome-wide association studies revealed associations between SMIM1 variants and Hb concentration and erythrocyte parameters^30^, particularly mean corpuscular Hb concentration (MCHC; P = 4.09 × 10^−96^) (Fig. 6m), suggesting that SMIM1 may play a more prominent role in maintaining iron homeostasis in humans than in mice during the steady state.

Together, our findings define a molecular function for the Vel blood group protein, identify SMIM1 as a previously unrecognized route for acquiring iron from extracellular Hb, and reveal an alternative, macrophage-independent pathway for Hb-derived iron recovery.

## Methods

### Plasmids

Lentiviral sgRNA vectors were generated based on lentiCRISPRv2 (Addgene 52961) or lentiGuide-Puro (Addgene 52963). The guide RNA sequences used in this study were ordered from IDT and are listed in Table S2. The SMIM1 sequence was purchased from Twist Biosciences and cloned into the doxycycline-inducible overexpression system pCW57.1 (Addgene 41393).

### Reagents

The following reagents were used throughout this study: Anisomycin (Sigma, A9789), Recombinant Human IFN-γ (peprotech, 300-02), Recombinant USP36 (R&D Systems, E-628) Hydrogen Peroxide, 30% (Fischer Scientific, H325-100), Dynasore (Selleckchem, S8047), Deferoxamine (Sigma, D9533), Bafilomycin A1 (Sigma, B1793), Hemoglobin human (Sigma, H7379), Hemoglobin bovine (Sigma, H2500), Ferritin human (Sigma, F6879), Transferrin human (Sigma, T8158), Haptoglobin from pooled human plasma (Sigma, H3536), 1-Oleoyl-2-palmitoyl-sn-glycero-3-phosphocholine (POPC) (Sigma/Merck, P4142), phenylhydrazine (PHZ; Sigma-Aldrich, 114715), human hemoglobin S (HbS, Sigma/Merck, H0392-25MG), human hemoglobin A2 (HbA_2,_ Sigma/Merck, H0266-5MG), human Hemoglobin A0 (HbA0, Sigma/Merck, H0267-25MG), Myoglobin from equine heart (Mb, Sigma/Merck, M1882-250MG), Hemin Chloride (Sigma/Merck, 3741-5GM), ferric (III) ammonium citrate (Sigma/Merck, F5879-100G).

### Cell culture

HAP1^31^, HepG2, and K562 cells (ATCC) were cultured in Iscove’s Modified Dulbecco’s Medium (IMDM, Gibco) supplemented with 10% heat-inactivated fetal calf serum (FCS) (Sigma), 1% penicillin-streptomycin (Thermo Fisher Scientific), and 1% L-Glutamine (Thermo Fisher Scientific), unless stated otherwise. HeLa and HEK293T cells (ATCC) were maintained in Dulbecco’s Modified Eagle Medium (DMEM) with 10% FCS, 1% penicillin-streptomycin, and 1% L-Glutamine (Thermo Fisher Scientific). All cell lines were grown at 37°C with 5% CO_2_ and were regularly tested for mycoplasma contamination.

### Lentiviral transductions

HEK293T cells (ATCC) were transfected with the packaging plasmids: 7.5 μg VSV-G (Addgene 8454), 3.75 μg psPAX-2 (Addgene 12260), and 10 μg of target vectors using TurboFectin 8.0 transfection reagent (TF81005, Origene). Viral supernatant was collected 48 hours post-transfection, filtered, and used for transduction in the presence of 8 μg/mL protamine sulfate. Transduced cells were selected with puromycin (HepG2: 2 μg/mL, HAP1: 1 μg/mL) or blasticidin (HepG2: 50 μg/mL, HAP1: 20 μg/mL) for 48 hours.

### FACS-based phenotypic screens

To generate libraries of mutagenized HAP1 cells with either gain- or loss-of-function constructs, gene-trap retrovirus for mutagenesis of HAP1 cells was produced using HEK293T cells, which were transfected with either an activating gene-trap vector containing a strong EF1a promoter or a disruptive vector, as described previously^32^. These constructs contain GFP to assess mutagenesis efficiency, along with Gag-pol, VSVg, and pAdv as packaging plasmids. The supernatant was harvested 48- and 72-hours post-transfection, filtered, concentrated using Amicon filters, and stored at 4°C. Retrovirus from both harvests was combined, supplemented with protamine sulfate (8 µg/ml), and used to infect 40 × 10^6^ wildtype HAP1 cells. After expansion, mutagenized cells were either treated or left untreated, as detailed in Table S3, depending on the molecular readout used. Cells were harvested with trypsin, resuspended in medium, pelleted, and washed with PBS.

For intracellular staining, approximately 4 billion cells were fixed in BD Fix Buffer I for 10 minutes at 37°C, washed twice with FACS buffer (PBS and 10% FCS), and permeabilized on ice with BD Perm Buffer III for 30 minutes. Cells were then washed once in FACS buffer and stained for 1 hour at room temperature with the indicated antibodies listed in Table S1 and S3, followed by secondary antibody staining for an additional hour, along with DAPI (1 µg/ml) to visualize G1 cells.

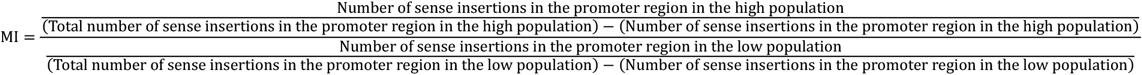

For surface staining, for example in the case of CD274 (see Table S3), after the cells were harvested with trypsin, resuspended in medium, pelleted, and washed with PBS, the cells were directly stained with primary antibody at room temperature for 1 hour in surface staining buffer (PBS, 0.5% FCS, 0.2% sodium azide). After two washes, secondary antibody staining was performed for an additional 1 hour in surface staining buffer. The cells were then washed twice with PBS, fixed with BD Fix Buffer I for 10 minutes at 37°C, and permeabilized on ice with BD Perm Buffer III for 15 minutes. After two washes, cells were stained for 1 hour with DAPI (1 µg/ml) to visualize G1 cells. After two final washes in FACS buffer, cells were filtered through a 40µm strainer (BD Falcon). At least 10^7^ cells per channel (~5% of the lowest and highest signal) were sorted using a BD FACSAria™ Fusion Cell Sorter. All antibodies used in the FACS-based gain-of-function genetic screens were previously validated in FACS-based loss-of-function genetic screens (Table S3), except where indicated otherwise. Genomic DNA was then isolated using a DNA mini kit (QIAGEN), and insertion sites were amplified using Linear Amplification Polymerase Chain Reaction (LAM-PCR), as described previously^8,9^.

### Insertion site mapping and data analysis

For the Gain-of-function genetics screens, after the next-generation sequencing was performed on the low and high sorted populations, the gene-trap insertion sites were identified as unique reads aligning unambiguously to the hg38 human genome using Bowtie^33^, allowing for a single mismatch. These aligned reads were mapped to hg38 protein-coding genes (RefSeq) to locate the insertion sites and determine their orientation with respect to the gene using IntersectBED^34^. This analysis was performed using the Phenosaurus pipeline (https://github.com/BrummelkampResearch/phenosaurus; last accessed 4 October 2026), and sense integrations in the promoter region, spanning from the 5′ UTR to −6 kbp upstream of the transcription start site (TSS), were analyzed. The number of unique activating insertions in each gene for the high population was compared to the low population using a two-sided Fisher’s exact test, with p-values adjusted for multiple testing using the Benjamini-Hochberg correction. A mutation index (MI) was calculated for each gene as the ratio of activating integrations, normalized by the total number of integrations across both high and low populations in the promoter region, using the following formula:

To prevent potential confounding effects caused by the possibility that the phenotype is mediated by a neighboring gene, we similarly mapped the insertions in the coding region of each gene (CDS-start to CDS-end). The number of unique activating insertions in each gene in the high population were then counted and compared with those in the low population, as above using a two-sided Fisher’s exact test followed by false discovery rate correction with the Benjamini-Hochberg method. The mutational index (MI) was calculated in the CDS regions similarly as above. For a gene to score as activating, it must score significantly in the promoter region but not in the CDS. If a gene scored significantly in both the promoter and CDS, we assessed the MI values. If the MI values in the promoter region had an opposing sign to the MI values in the CDS region, the gene was retained as a hit (rate-limiting factor). However, if they had the same sign, the hit was excluded, as the phenotype is more likely to be caused by a neighboring gene. For the loss-of-function genetic screens, we used the same approach described previously^8,9^ and explained above, taking into consideration the sense unique insertions within a gene (5′ UTR + CDS). Integrations within genomic regions associated with overlapping genes were excluded, along with insertions in the 3′ untranslated region (3’UTR), where the gene-trap cassette is less effective in disrupting gene function. The FDR-corrected *p*-value and the MI were calculated as described above.

### Flow cytometry analysis

For intracellular staining, 3 × 10^6^ HAP1 cells were seeded in 10-cm dishes and treated as indicated. Cells were first trypsinized, washed once with PBS, and fixed with BD Phosflow Fix Buffer I for 10 minutes at 37°C. Next, cells were washed once with PBS and permeabilized using BD Perm Buffer III for 30 minutes on ice. Samples were then washed once with PBS and blocked with FACS buffer (10% FCS in PBS) for 1 hour before incubation with primary antibodies in FACS buffer (Table S1) for an additional hour at room temperature. After 2 washes with PBS, secondary antibodies (Alexa-647/488, Invitrogen) and DAPI (1 μg/mL) were added for 1 hour. Finally, cells were washed twice with PBS, resuspended in FACS buffer, and passed through a 40µm strainer before analysis using a BD LSR Fortessa Cell Analyzer. For surface staining, cells were stained before fixation with the primary and secondary antibodies either at RT or on ice for 1 hour using surface staining buffer (PBS, 0.5% FCS, 0.2% sodium azide). The cells were then fixed, permeabilized, stained with DAPI, and analyzed by flow cytometry as described above. Data analysis was carried out using FlowJo software version 10.9.0.

### Protein extracts and immunoblotting

Cell pellets were lysed in RIPA buffer containing protease inhibitors (Roche). Following sonication, protein concentrations were measured using the BCA Protein assay Kit (Pierce, Thermo Fisher Scientific). Equal amounts of protein were separated by SDS-PAGE and transferred to nitrocellulose membranes (Bio-Rad). Protein detection was performed using primary antibodies listed in Table S1, following standard protocols. Alternatively, SDS sample buffer (0.25 M Tris-HCl pH 6.8, 6% SDS, 30% glycerol, 100 mM DTT, and bromophenol blue) was added directly to PBS-washed cells, and the samples were analyzed by SDS-PAGE as described above. Immunoblots were blocked with 5% BSA (Sigma, A3059) in TBS-Tween.

### Immunofluorescence

Cells were seeded in 8-well chambers (Thermo Fisher Scientific) and treated as described in the figure legends. To stain the plasma membrane, we used CellMask (C37608, Thermo Fisher), and to visualize hemoglobin (Hb), we labeled Hb using Alexa Fluor™ 647 NHS Ester (Thermo Fisher Scientific, A20006). Briefly, 2 mg/ml of Hb dissolved in 500 µl of PBS was mixed with 500 µl of 0.1 mg/ml Alexa Fluor™ 647 NHS Ester dissolved in PBS and incubated for 30 minutes at room temperature. The labeled Hb-AF647 was then purified using a desalting column and concentrated with Amicon Ultra-0.5 ml centrifugal filters (Merck Millipore, UFC500396). Cells were fixed for 10 min using either 4% alcohol-free formaldehyde or ice-cold 100% methanol (for PAR staining). After fixation, cells were permeabilized for 1 h in IF buffer (0.2% Triton X-100, 10% FCS in PBS). Following permeabilization, slides were incubated with primary antibodies diluted in IF buffer for 1 hour at room temperature. The cells were then washed three times with PBS and incubated with secondary antibodies (Invitrogen) and DAPI (1 µg/ml, Sigma) for an additional hour. After a final series of washes, slides were mounted using Aqua-Poly/Mount mounting medium (Polysciences). Images were acquired with a ZEISS LSM 980 Airyscan microscope.

### Proteomics

The samples were reconstituted in 1x S-Trap Lysis buffer (5% SDS, 50 mM TEAB pH 8.5). Protein content was determined using a BCA Protein Assay Kit (Pierce), after which 30 µg-aliquots of protein were reduced with 5 mM DTT and alkylated with 20 mM chloroacetamide by heating for 10 min at 95^°^ C on a thermoshaker. Proteins were acidified and digested overnight at 37^°^ C with trypsin (Sigma-Aldrich; 3 µg per sample) on S-Trap Micro spin columns following the manufacturer’s instructions (ProtiFi, NY, USA). Peptides were eluted, vacuum-dried and stored at −80°C until LC-MS/MS analysis. Peptides were analyzed by data-dependent acquisition (DDA) on an Orbitrap Exploris 480 Mass spectrometer connected to an Evosep One nano-LC system (Evosep Biotechnology, Odense, Denmark). Prior to LC separation with the Evosep One, peptides were reconstituted in 0.1% formic acid and 0.5µg was loaded on Evotip Pure™ tips (Evosep). Peptides were then eluted and separated using the pre-programmed (88 min gradient) on an EV1137 column (Evosep) with an EV1086 emitter (Evosep). Nanospray was achieved using the Easy-Spray NG Ion Source (Thermo Scientific) with spray voltage set to 1850V. The Exploris 480 was operated in DDA Cycle Time mode with 1 sec. Duty cycle and full MS scans being collected in the Orbitrap analyzer with 60,000 resolution at m/z 200 over a 375-1500 m/z range. Default charge state was set to 2+, the AGC target was set to “standard” for both MS1 and MS2 and for both scan types. The maximum injection time mode was set to “auto”. Monoisotopic peak determination was set to “peptide” and dynamic exclusion was set to 20 sec. For MS2, a normalized HCD collision energy of 30% was applied to precursors with a 2+ to 6+ charge state meeting a 5e^4^ intensity threshold filter criterium. Precursors were isolated in the Quadrupole analyzer with a 1.2 m/z isolation window and MS2 spectra were acquired at 15,000 resolution in the Orbitrap. RAW files were analyzed by label-free quantitation (LFQ) with Proteome Discoverer (PD, Thermo Scientific, version 3.1.0.638) using standard settings. Fragment spectra were searched against the Swissprot reviewed bovine database (release 2024_04) using SequestHT with Percolator. The maximum allowed precursor mass tolerance was

50 ppm and 0.06 Da for fragment ion masses. Trypsin was specified as proteolytic specificity allowing a maximum of 2 miscleavages; acetylation (protein N-term) and oxidation (M) were selected as variable modifications, whereas carbamidomethylation (C) was selected as fixed modification. False discovery rates for peptide and protein identification were set to 1% and as additional filter Sequest HT XCorr > 1 was set. Peptide spectrum match (PSM) numbers were used as a basis for protein abundance rank calculations in the 40% APS FCS fractions.

### SMIM1 peptides

SMIM1 peptides were ordered from Biomatik and corresponded to the C-terminal 15 amino acids of the wild-type SMIM1 extracellular domain (SMIM1-ECD-WT: FILGYLTGYYVHKCK) or an alanine-mutated HKCK motif variant (SMIM1-ECD-4A: FILGYLTGYYVAAAA). The peptides were used for FACS, immunofluorescence, liposome, and isothermal titration calorimetry (ITC) experiments. Similar N-terminally biotinylated peptides were used for surface plasmon resonance (SPR) assays.

### Surface Plasmon Resonance (SPR)

The Biacore 1S+ instrument (Cytiva) was used to investigate the interaction between Hb, transferrin, Mb, HbA0 or HbA2 and SMIM1. N-terminally biotinylated SMIM1-ECD-WT (Biotin-FILGYLTGYYVHKCK) or SMIM1-ECD-4A (Biotin-FILGYLTGYYVAAAA) peptides were immobilized on the surfaces of two separate flow cells of an SA Series S Sensor Chip (Cytiva). Six increasing concentrations of Hb, transferrin, Mb, HbA0, or HbA2, in 2-fold steps, were injected over the chip surface in a buffer containing PBS and 0.05% Tween using single-cycle kinetics. An empty flow cell was used as the binding reference. At the end of each injection cycle, 0.05% SDS was applied as the regeneration solution. Data analysis was performed using Biacore Evaluation Software (Cytiva), and final figures were prepared with GraphPad Prism (GraphPad Software LLC).

### Isothermal Titration Calorimetry (ITC)

The binding of the SMIM1-ECD-WT (FILGYLTGYYVHKCK) to Hb was assessed using a MicroCal PEAQ-ITC instrument (Malvern Panalytical Ltd). Hb in PBS buffer at a concentration of 75 µM was loaded into the instrument’s cell, and SMIM1 in the same buffer at 3 mM was titrated into the cell. The titration protocol included an initial injection of 0.5 µl, followed by seven injections of 5 µl each, with a delay of 180 seconds between injections. The titration was performed four times, with only the excess mixture from the previous titration being removed from the cell before the next experiment. Data from all four titrations were combined using MicroCal Concat software (Malvern Panalytical Ltd) prior to analysis. A similar set of titrations, where SMIM1 was titrated into the PBS buffer in the cell, served as the control. Data analysis was carried out using the MicroCal PEAQ-ITC Analysis software (Malvern Panalytical Ltd).

### DUB assay

Nucleosome reconstitution and its PRC1-dependent ubiquitination (NCP K118/119) were performed as described previously^35^. The ubiquitinated NCP K118/119 was diluted to 1 μM using the DUB buffer (25 mM HEPES, 150 mM NaCl, 1 mM DTT, pH 7.5). BAP1/ASXL1 was used as a positive control and purified as described previously^35^, then diluted in DUB buffer to 200 nM. Recombinant USP36, purchased from R&D Systems (E-628), was diluted to 200 nM, 100 nM, 50 nM, and 25 nM in DUB buffer. Next, the prepared BAP1/ASXL1 and USP36 concentrations were added to the pre-diluted ubiquitinated NCP K118/119 in a 1:1 ratio. The samples were incubated at 37°C for 30 minutes, and the reaction was quenched by boiling with SDS buffer for 5 minutes. The samples were then loaded onto an SDS-PAGE gel, run for 45 minutes, and stained with Coomassie.

### Liposomes preparation

Liposomes were prepared as previously described with modifications^36^. Briefly, 1-Oleoyl-2-palmitoyl-sn-glycero-3-phosphocholine (POPC) (Sigma/Merck, P4142) was dissolved in a sufficient volume of chloroform to allow the lipid solution to evenly cover the entire bottom surface of a glass beaker. After 15 min of mixing, the chloroform was evaporated under a gentle argon flow with continuous mixing to generate a thin lipid film, which was subsequently hydrated with PBS under continuous mixing for 1 h and then incubated overnight at 4°C. The liposomal suspension was sonicated for 1 min (130 W, 40 kHz) to reduce particle size and minimize aggregation, followed by 12 cycles of extrusion through a 0.2 µm polycarbonate membrane. Liposomes were then purified by ultracentrifugation at 150,000 × g for 90 min at 4°C. The liposomal pellet was collected and resuspended in PBS for downstream applications.

### Generation and maintenance of mouse strains

C57BL/6N-Smim1^em1(IMPC)Wtsi^/WtsiIeg mice (RRID: IMSR_EM:11219) were originally generated at the Wellcome Trust Sanger Institute by CRISPR-mediated zygote microinjection. To establish the strain for this study, cryopreserved sperm was obtained from the European Mouse Mutant Archive (EMMA), and mice were rederived in-house. Genotyping was performed by standard PCR using the following primers, as indicated in Extended Data Fig.13A-B. WT primers (P1): Smim1_R_wt: TCCCCATCTCCAGTACACAGGGTA and Smim1_F_U: CCTGCACAGAAAGATTTGAGGCCC or mutant primers (P2): Smim1_R_ko: CAGAGGCCCCAGTTGTAAATGCAG and Smim1_F_U: CCTGCACAGAAAGATTTGAGGCCC.

The mouse colony was housed in a certified animal facility under specific pathogen-free conditions in temperature- and humidity-controlled rooms maintained at 21°C and 55% relative humidity. Animals were housed in individually ventilated cages under a 12 h light/12 h dark cycle with food and water provided ad libitum. All animal experiments were approved by the Animal Ethics Committee of the Netherlands Cancer Institute and performed in accordance with institutional, national, and European guidelines for animal care and use.

### Phenylhydrazine-induced anemia in mice

Female and male 8-10 weeks old wild-type (*Smim1*^*+/+*^) and SMIM1 knockout (*Smim1*^*−/−*^) mice were injected subcutaneously with phenylhydrazine (PHZ; Sigma-Aldrich, 114715) at a dose of 100 mg/kg body weight or with vehicle control (PBS). Animals were monitored twice daily for body weight and general welfare. Mice were euthanized by cervical dislocation 72 h after PHZ treatment, and the tibiae and femur were collected for subsequent analyses.

### Bone marrow isolation and flow cytometry analysis

Bone marrow cells were harvested from the tibiae and femur of 8-10 week-old wild-type (Smim1^+/+^) and SMIM1 knockout (Smim1^−/−^) mice challenged with either PHZ or vehicle control (PBS) by flushing the bones with cold PBS using a 27G needle and a 10 mL syringe. Cells were resuspended by gentle pipetting and passed through a 70 µm cell strainer. The cells were then centrifuged at 300 × g for 10 min at 4°C and resuspended in FACS buffer (PBS supplemented with 10% FCS).

For phenotypic analysis by flow cytometry, cells were counted, and 10^7^ cells were resuspended and stained in 300 µL of surface staining buffer (PBS + 0.5% FCS + 0.2% sodium azide) containing a 1:100 dilution of the following antibody cocktail: Spark PLUS UV395™ anti-mouse CD45 antibody (BioLegend, 103192), PE anti-mouse TER-119 (TER-119) (Cytek, 50-5921-U100), PE/Cyanine7 anti-mouse CD71 antibody (BioLegend, 113812), FITC anti-mouse CD117 (c-Kit) antibody (BioLegend, 105806), and Alexa Fluor® 647 anti-mouse/human CD44 antibody (BioLegend, 103018). Cells were incubated on ice in the dark for 30 min. If PE/Cyanine7 anti-mouse CD71 antibody was included in the staining panel, True-Stain Monocyte Blocker (BioLegend, 426102) was used to block non-specific PE/Cyanine7 binding. Cells were subsequently washed twice with 1 mL of FACS buffer and passed through a 40 µm cell strainer. DAPI was added immediately before acquisition to distinguish live and dead cells. Data acquisition was performed using a BD Symphony A5 flow cytometer with DiVA software (BD Biosciences), and data analysis was carried out using FlowJo software version 10.9.0.

## Data availability

Raw data generated from the FACS-based haploid genetic screen have been deposited in the European Nucleotide Archive under accession PRJEB120955. The corresponding screening results are also available through an interactive web interface at https://phenosaurus.nki.nl.

## Acknowledgements

We would like to thank the members of the Brummelkamp group for their valuable discussions and Patrick Celie from the NKI Protein Facility for his assistance with hemoglobin labeling. We thank John Strouboulis from the Faculty of Life Sciences & Medicine, King’s College London, for critically reading the manuscript. We also thank Titia Sixma for valuable discussions and for reading the manuscript. We extend our gratitude to the NKI Flow Cytometry Facility, particularly Martijn van Baalen, as well as the Genomics Core Facility, and Proteomics Facility for their technical support. Schematic illustrations were created using and adapted from BioRender.com.

Research at the Netherlands Cancer Institute is funded by institutional grants from the Dutch Cancer Society and the Dutch Ministry of Health, Welfare, and Sport. The T.R.B. laboratory receives support from the NWO Vici grant (016.Vici.170.033), the Oncode Institute and European Research Council (ERC) advanced grant (101142218). A.M. is funded by an EMBO Long-Term Fellowship (ALTF 158-2018). JJ is supported by the Oncode Institute, which is partially funded by the Dutch Cancer Society and the Mark Foundation for Cancer Research.

## Author contributions

This study was conceptualized by A.M. and T.R.B.

A.M. performed, analyzed, and visualized the experiments. D.B. assisted with experiments related to SMIM1. A.M. performed all the genetic screens and analyzed the genetic-screen data with support from M.L.H., G.H.P., and L.W. A.M. and A.F. performed surface plasmon resonance (SPR) and isothermal titration calorimetry (ITC) assays.

A.M. and S.S. performed the DUB assays. A.M., C.L., S.C.M., and B.K. performed the *in vivo* intervention study under the supervision of J.J. and H.A.M. M.D. organized the mouse colonies. O.B.B. performed and analyzed the proteomics experiments. A.M. and T.R.B. wrote the manuscript with input from all authors. The study was supervised by A.M. and T.R.B.

---

**Extended Data Figure 1.**
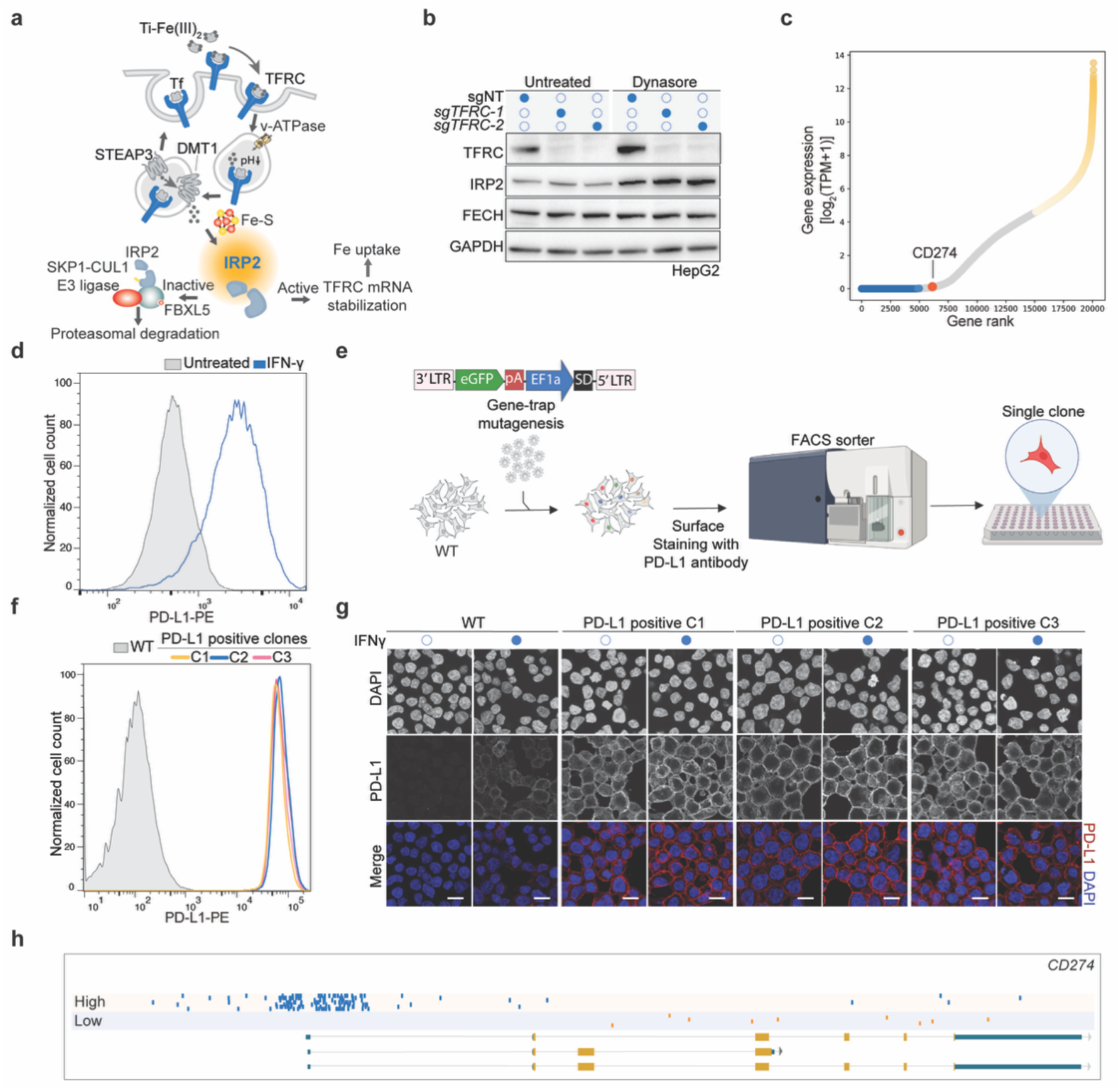
Designing FACS-based gain-of-function genetic screening using retroviral integration in human cells. (**a**) IRP2 acts as a central sensor for iron uptake mediated by endocytosis of diferric transferrin through the TFRC pathway. (**b**) sgNT or sgTFRC HepG2 cells were left untreated or treated with dynasore (10 µM) for 24 hours. The whole-cell extracts were subjected to immunoblot analysis using the indicated antibodies, with GAPDH used as a loading control. (**c**) Ranking plot of gene expression in wild-type HAP1 cells. Genes were ranked based on log_2_ (TPM + 1), where TPM represents transcripts per million, using published RNA-sequencing data^37^. The 5,000 highest- and lowest-expressed genes are highlighted in yellow and blue, respectively. CD274, encoding PD-L1, is indicated by a red dot. (**d**) Flow cytometry analysis of PD-L1 cell surface expression in wild type HAP1 cells treated with 0.5 ng/ml IFNγ for 24 hours or left untreated. (**e**) Schematic overview of the experimental setup for validating the gain-of-function strategy. Wild type HAP1 cells were transduced with a retroviral activating gene-trap vector containing the EF1a promoter. Random insertions of EF1a across the genome created a pool of cells with different genotypes, which were stained for surface PD-L1, followed by single-cell sorting of cells with high PD-L1 expression. (**f-g**) PD-L1 surface expression levels were measured in three independent clones derived from (e) after FACS-based sorting using PD-L1 as a marker, either by flow cytometry (f) or cell surface immunofluorescence (g). Scale bar, 10 μm. (**h**) Enrichment of activating gene-trap insertions in the *CD274* (PD-L1) locus in cells selected for PD-L1 expression. Blue and yellow dots represent sense insertions in the high and low populations, respectively.

**Extended Data Figure 2.**
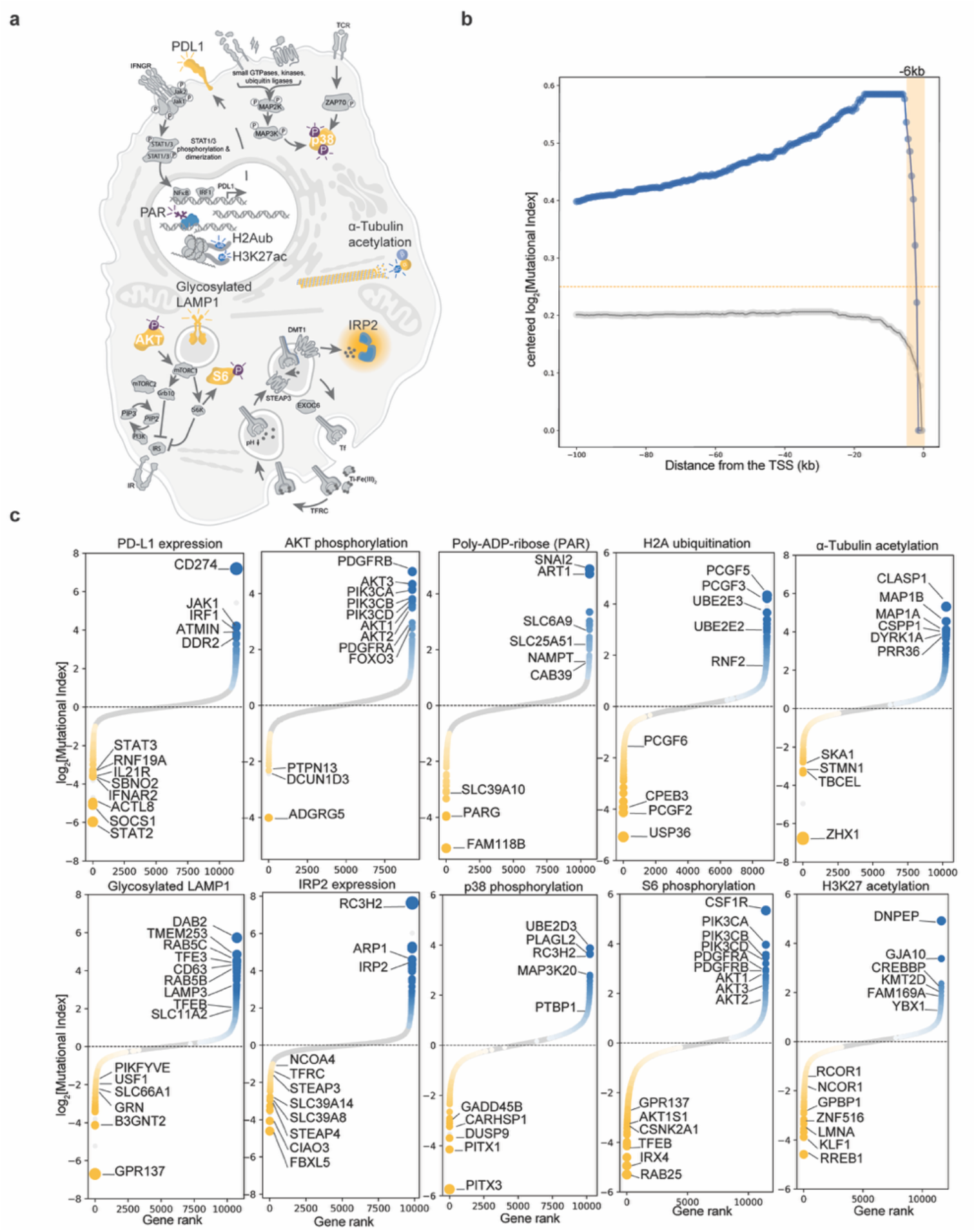
Gain-of-function FACS-based genetic screens across diverse biological processes. (**a**) Overview of the molecular readouts used for the FACS-based haploid genetic screens using gain- and loss-of-function mutagens. (**b**) Impact of sense insertions varying the distance from the TSS by steps of 1 kb on the phenotype of PD-L1 expression. Blue line represents the median absolute mutational index of genes where at least one of the assessed intervals was significantly enriched for sense insertions (*p*-value ≤ 0.05, Log_2_ MI > 1), while the gray line shows the same but for non-significant genes. (**c**) Hits identified using Gain-of-function genetic screens conducted using the indicated antibodies. For each gene (represented by dots), the mutation index (MI) is plotted on the y-axis. The data is displayed as a ranking plot according to MI values. Genes significantly enriched in either channel (two-sided Fisher’s exact test; FDR-corrected p-value ≤ 0.05; total number of unique insertions > 50) are color-coded, and selected regulators are labelled

**Extended Data Figure 3.**
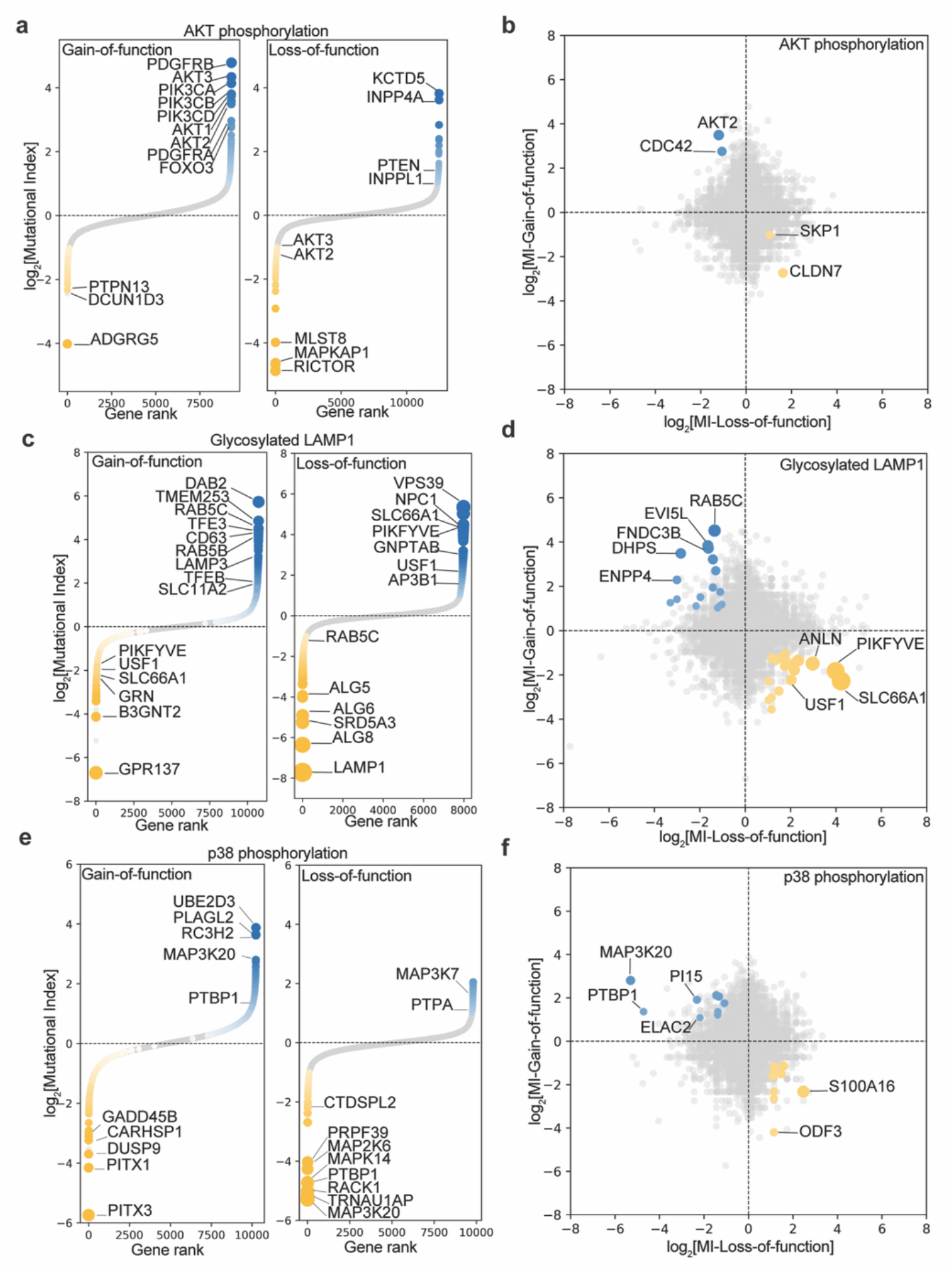
The gain-of-function approach identifies rate-limiting factors across a range of molecular readouts. (**a-f**) Examples of gain- and loss-of-function genetic screens were performed using AKT phosphorylation, glycosylated LAMP1, and p38 phosphorylation. The results are shown as ranking plots based on mutation index (MI) values. Genes significantly enriched in either channel (two-sided Fisher’s exact test; FDR-corrected p-value ≤ 0.05; total number of unique insertions > 50) are color-coded, with top regulators labeled. Ranking plots are paired with scatter plots comparing MI values for the loss-of-function screen (x-axis) to the gain-of-function (y-axis), with selected inversely correlated genes are labeled.

**Extended Data Figure 4.**
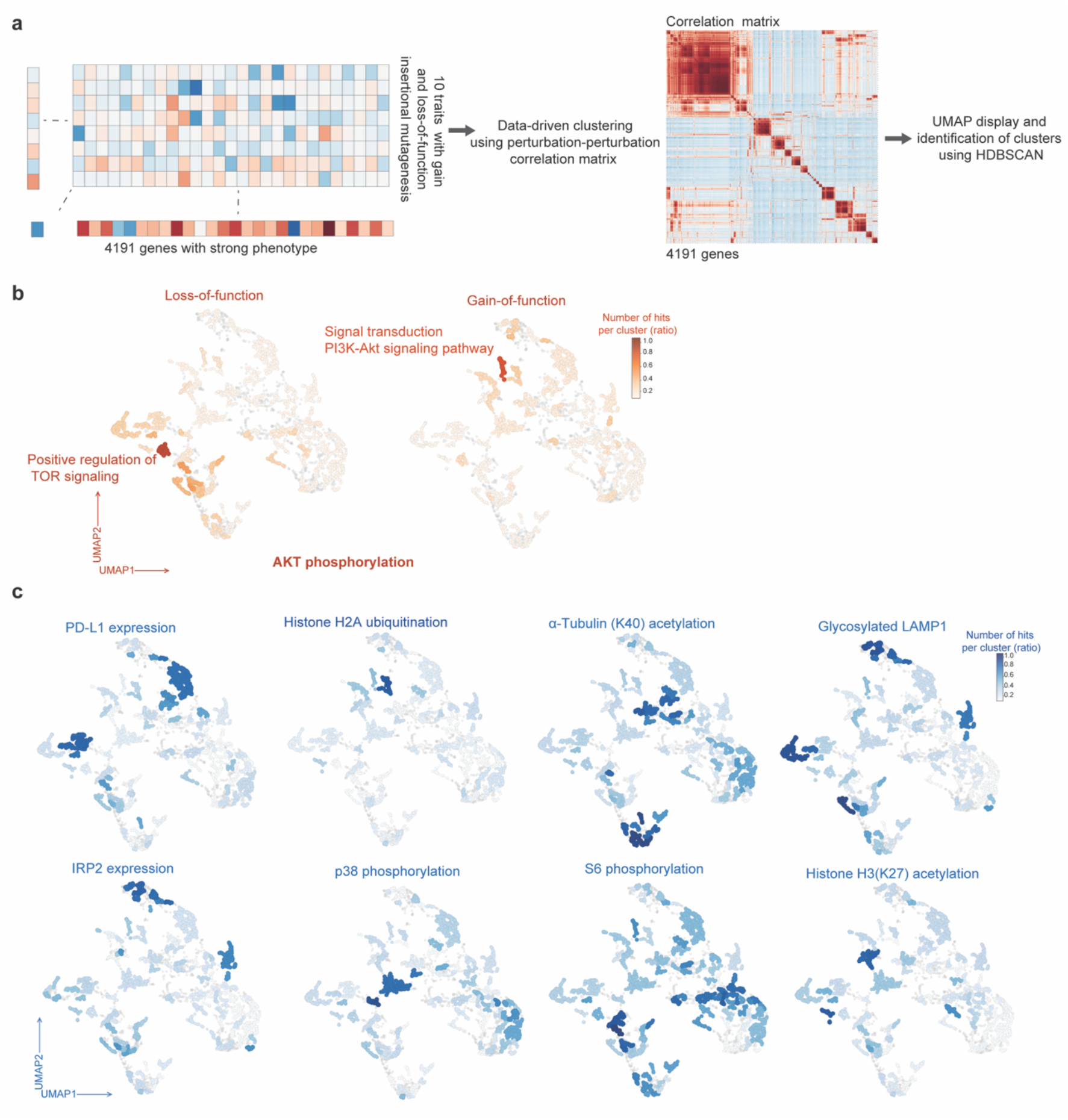
Exploring genotype-phenotype relationships using complementary gain- and loss-of-function genetic approaches. **(a)** Analysis workflow overview. 20 phenotype parameters were aggregated into profiles for each of the 4192 genes. The relationships between profiles were visualized using UMAP and the clusters were identified using HDBSCAN. **(b)** Significant hits (FDR-corrected *p*-value ≤ 0.05, and Log_2_[MI]> 1 or Log_2_[MI] < −1) from the AKT phosphorylation screen, either using loss-of-function or gain-of-function are displayed on the UMAP. The number of hits in each HDBSCAN cluster is represented by color contrast.(**c**) Significant hits from the indicated molecular readout screens, conducted using both loss- and gain-of-function genetic approaches, are displayed on the UMAP and colored as indicated in (b).

**Extended Data Figure 5.**
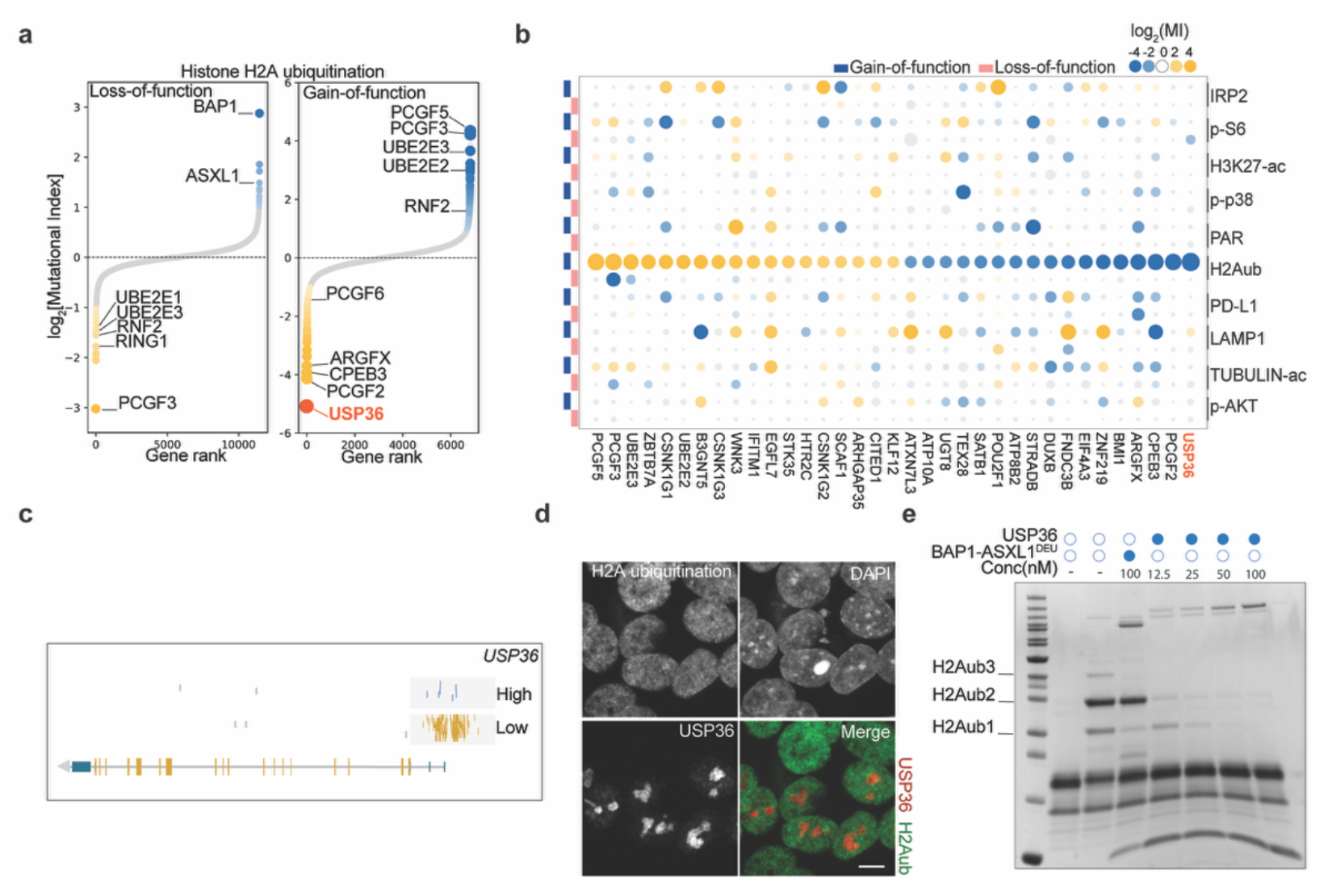
USP36 ensures the deubiquitination of H2A. (**a**) H2A ubiquitination screens were performed using both loss- and gain-of-function haploid genetic approaches. The results are presented as ranking plots based on mutation index (MI) values. Genes significantly enriched in either channel (two-sided Fisher’s exact test; FDR-corrected p-value ≤ 0.05; total number of unique insertions > 50) are color-coded and top regulators are labeled. USP36 is highlighted with a red dot. (**b**) Bubble plot displaying the top-scoring hits of the gain-of-function H2A ubiquitination screen (FDR-corrected *p*-value ≤ 0.05 and Log_2_[MI] > 2 or Log_2_[MI] < −2) across various FACS-based loss- and gain-of-function genetic screens with different cellular readouts. Regulators that increase the readout signal are shown in yellow, while those that decrease it are shown in blue. The intensity of the colors and the size of the bubbles represent Log_2_[MI] values, with non-significant hits highlighted in grey. The hits are ranked based on the Log_2_[MI] values from the gain-of-function H2A ubiquitination genetic screen. (**c**) Activating sense insertions identified in the USP36 locus are shown in the high channel (High, blue) compared to the low channel (Low, yellow) in the gain-of-function genetic screen using H2A ubiquitination as a readout. (**d**) Immunofluorescence staining for H2A ubiquitination (K119) and HA-USP36 in wild-type HAP1 cells. Scale bar, 5 μm. (**e**) Deubiquitinase activity assay of USP36 recombinant protein incubated with 0.5 µM H2A K119 ubiquitinated oligonucleosomes at the indicated concentrations for 30 minutes. The BAP1/ASXL1^DEU^ complex was used as a positive control. The reactions were visualized on SDS-PAGE and stained with Coomassie.

**Extended Data Figure 6.**
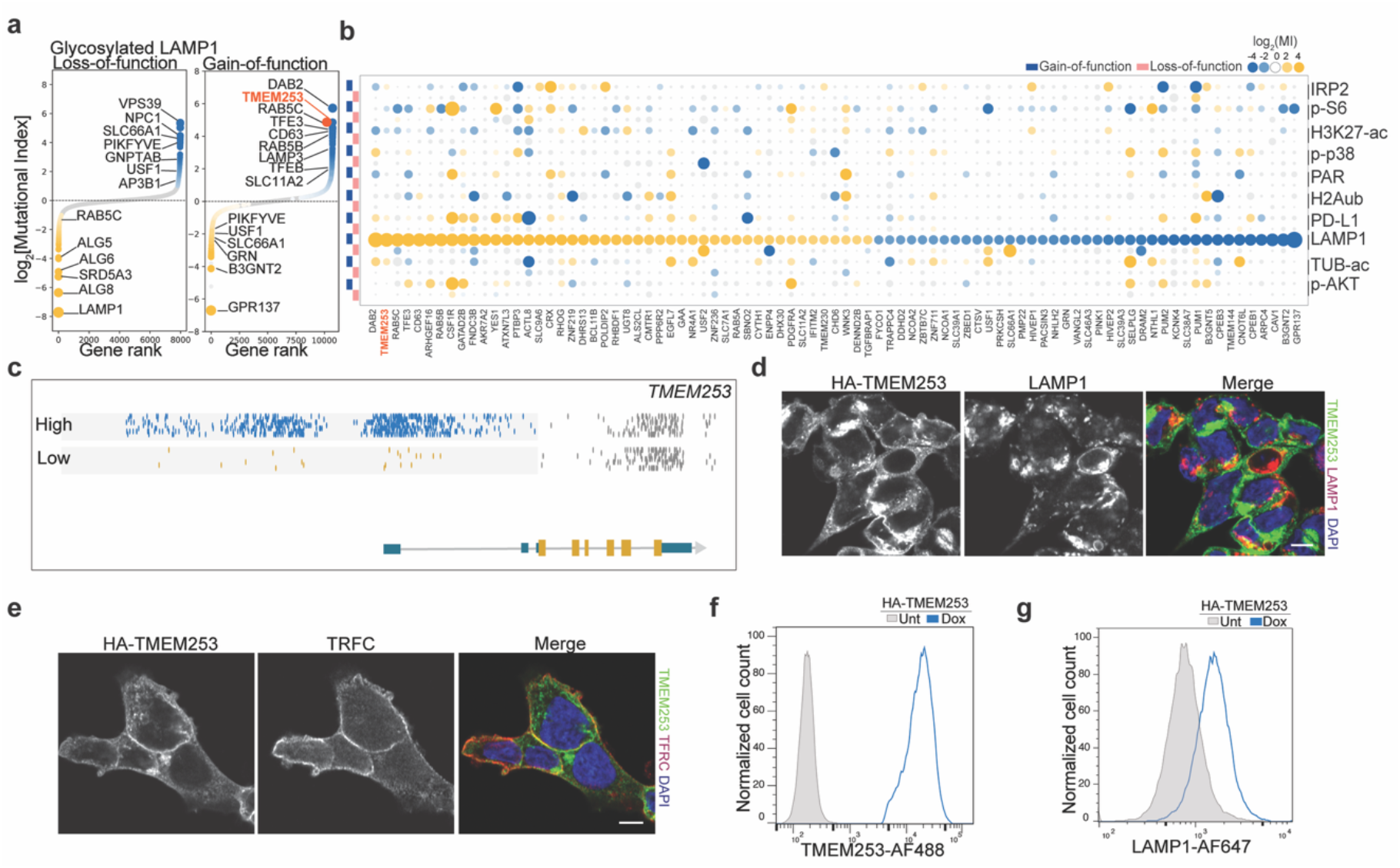
TMEM253 regulates LAMP1 protein abundance. (**a**) LAMP1 abundance screens conducted using both loss- and gain-of-function haploid genetic approaches. The data is presented as ranking plots based on mutation index (MI) values. Genes significantly enriched in either channel (two-sided Fisher’s exact test; FDR-corrected p-value ≤ 0.05; total number of unique insertions > 50) are color-coded and top regulators are labeled. TMEM253 is shown in red. (**b**) Bubble plot showing the top-scoring hits (FDR-corrected *p*-value ≤ 0.05 and Log_2_[MI] > 2 or Log_2_[MI] < −2) across various FACS-based loss- and gain-of-function genetic screens, using different biological readouts as indicated. Regulators that increase the readout signal are colored yellow, while those that decrease the signal are in blue. The contrast of the colors and the size of the bubbles correspond to Log_2_[MI]. Non-significant hits are highlighted in grey. (**c**) Activating sense insertions identified in the TMEM253 locus in the high channel (High, blue) compared low channel (Low, yellow) in the gain-of-function genetic screen using glycosylated LAMP1 as a readout. (**d-e**) Immunofluorescence staining displaying colocalization of HA-TMEM253 with LAMP1 (d) or TFRC (g) in wild-type HAP1 cells. Scale bar, 5 μm. (**f-g**) Wild-type HAP1 cells with doxycycline-inducible TMEM253 (N-terminus HA tag) were either induced with doxycycline for 48 hours or left untreated as a control. These cells were then fixed, permeabilized, and stained with HA-TMEM253 (f) and LAMP1 (g) antibodies. the cells were then subjected to flow cytometry analysis.

**Extended Data Figure 7.**
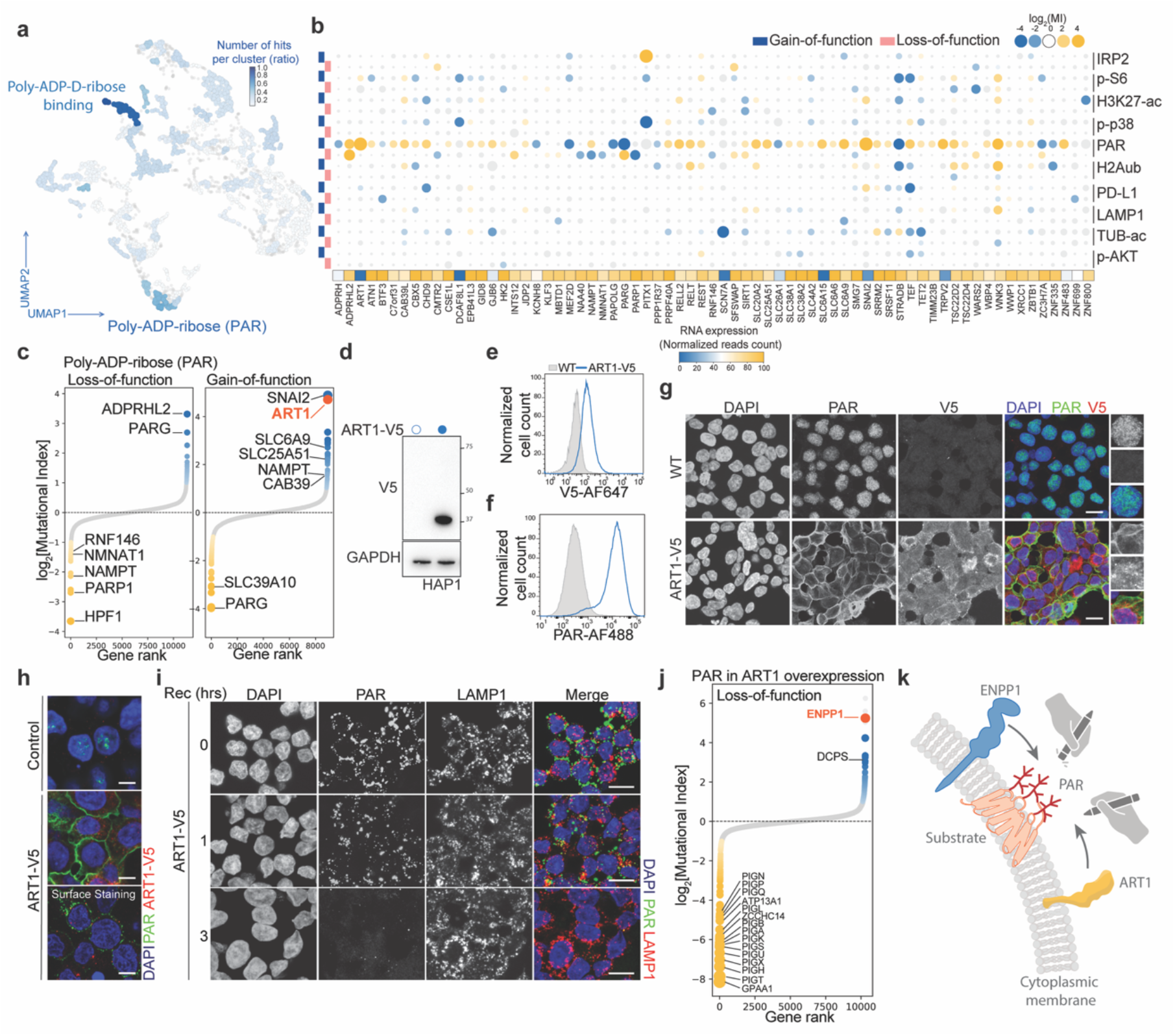
Complementary gain- and loss-of-function screens to study poly-ADP ribosylation. **(a)** Significant hits from the PAR screens conducted using both loss- and gain-of-function genetics are displayed on the UMAP with the number of hits in each HDBSCAN cluster represented by color intensity. (**b**) Bubble plot depicting the most represented HDBSCAN cluster in the PAR screens. The heatmap (squares) indicates RNA expression levels of the indicated genes in HAP1 cells. (**c**) PAR screens performed using both loss- and gain-of-function haploid genetic approaches. The results are displayed as ranking plots based on mutation index (MI) values. Genes significantly enriched in either channel (two-sided Fisher’s exact test; FDR-corrected p-value ≤ 0.05; total number of unique insertions > 50) are color-coded and selected regulators are labeled. (**d**) Whole-cell extracts from V5-tagged ART1-expressing cells and control wild-type HAP1 cells, which were subjected to immunoblot analysis using a V5 antibody, with GAPDH as a loading control. (**e**-**f**) Wild-type HAP1 cells expressing V5-tagged ART1, or control cells were fixed, permeabilized, and stained with V5 (e) and PAR (f) antibodies, and DNA was stained using DAPI. The cells were then subjected to flow cytometry analysis. (**g**) Wild-type HAP1 cells expressing V5-tagged ART1 or control cells were fixed, permeabilized, and stained for PAR and V5-ART1. Scale bar, 10 μm. (**h**) The same cell lines used in (g) were either fixed and permeabilized (upper and middle panels) or surface-stained for PAR (lower panel). Scale bar, 5 μm. (**i**) Wild-type HAP1 cells expressing V5-tagged ART1 were surface-stained in live cells using an antibody directed against PAR. Cells were then fixed at the indicated recovery time points, followed by permeabilization and staining with an antibody directed against LAMP1. DAPI was used to stain DNA. Scale bar, 10 μm. (**j**) Loss-of-function genetic screen in ART1-V5 expressing cells using PAR surface staining. The mutagenized cell population was surface stained using antibody directed against PAR, followed by fixation, permeabilization, and staining with antibody directed against the V5 epitope and DAPI. G1-phase cells expressing the same amount of ART1 with the highest and lowest 5% PAR signals were sorted, and the gene-trap insertion sites were mapped using next generation sequencing. The data is presented as ranking plots based on mutation index (MI) values. Genes significantly enriched in either channel (two-sided Fisher’s exact test; FDR-corrected p-value ≤ 0.05; total number of unique insertions > 50) are color-coded and selected regulators are labeled. (**k**) Proposed model of the regulation of PAR at the plasma membrane through the ART1-ENPP1 axis.

**Extended Data Figure 8.**
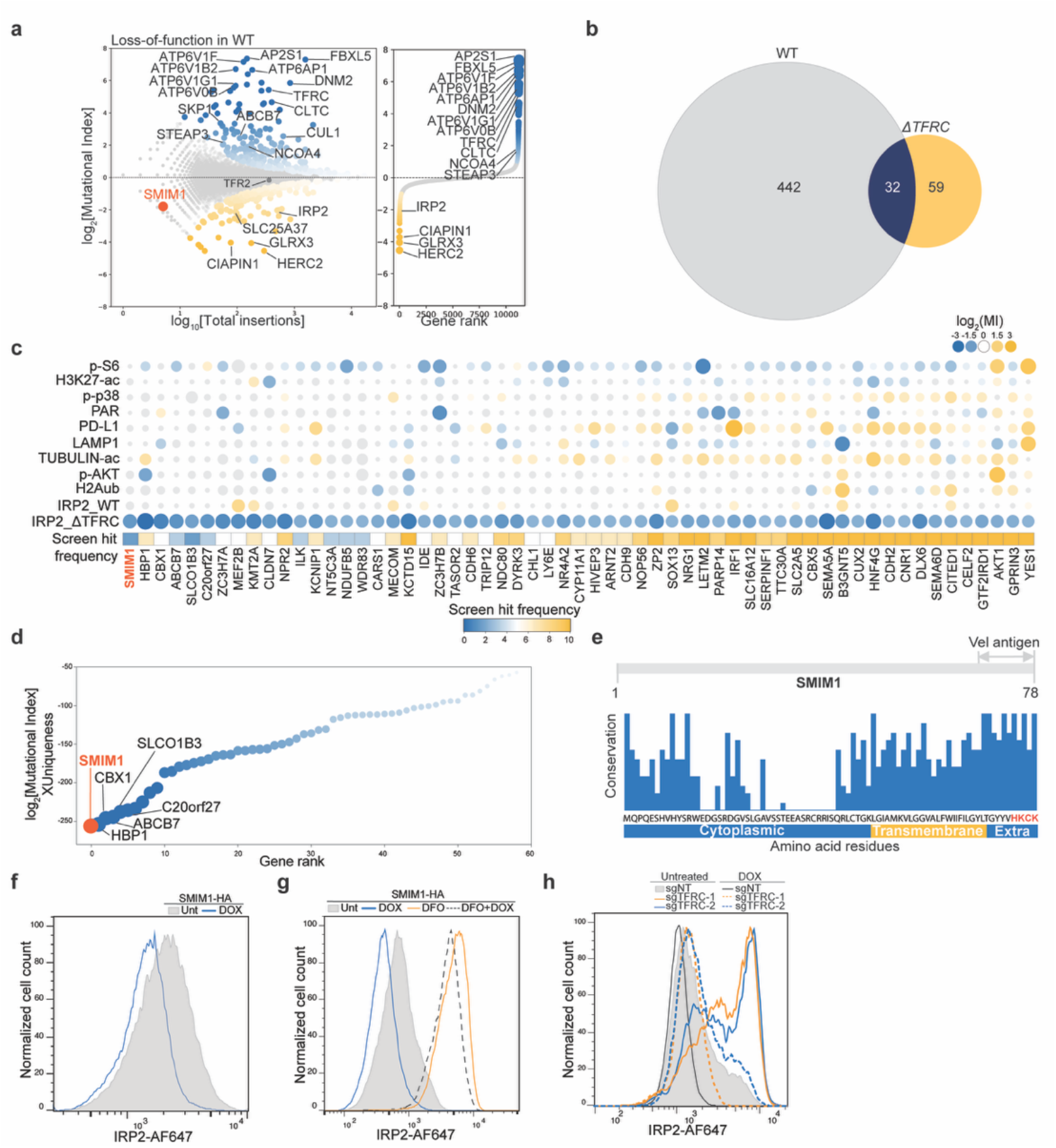
SMIM1 modulates iron metabolism. (**a**) A fishtail plot of the FACS-based loss-of-function genetic screen using IRP2 as a readout in wild type HAP1 cells is shown. The y-axis displays the mutational index (MI) for each gene, and the x-axis indicates the total number of insertions per gene. The ranking plot represents in x-axis the gene rank based on the mutational index. (**b**) Venn diagram of the factors that strongly decrease IRP2 levels (FDR-corrected *p*-value ≤ 0.05 and Log_2_[MI] < −1) identified using the gain-of-function genetic screens in wild type or *TFRC* knockout HAP1 cells. (**c)** A bubble plot displaying the 59 hits that caused a significant reduction in IRP2 protein levels specifically in TFRC knockout cells, but not in wild-type HAP1 cells shown in (b). The hits are ranked according to their impact on IRP2 levels and the uniqueness of their scores across 10 distinct molecular readouts from the gain-of-function screens. (**d**) The hits from (c) are displayed as a ranking plot. Top outliers are labeled, and SMIM1 is highlighted in red. (**e**) Schematic depiction of the SMIM1 cytoplasmic, transmembrane, and extracellular domains (Vel antigen) and the degree of SMIM1 conservation across vertebrates. (**f**) Wild type HAP1 cells with doxycycline-inducible SMIM1 (C-terminal HA tag) were induced with 2 µg/ml doxycycline for 48 hours or left untreated as a control. The cells were then stained for IRP2 and analyzed by flow cytometry. (**g**) Wild type HAP1 cells expressing doxycycline-inducible SMIM1 (C-terminal HA tag) were treated with 2 µg/ml doxycycline for 24 hours to induce SMIM1 expression or left untreated as a control. Cells were then either left untreated or exposed to 10 µM deferoxamine (DFO) for an additional 24 hours, followed by IRP2 staining and flow cytometric analysis. (**h**) Data from Figure 3g is presented in the same flow cytometry histogram, showing IRP2 levels in the indicated conditions.

**Extended Data Figure 9.**
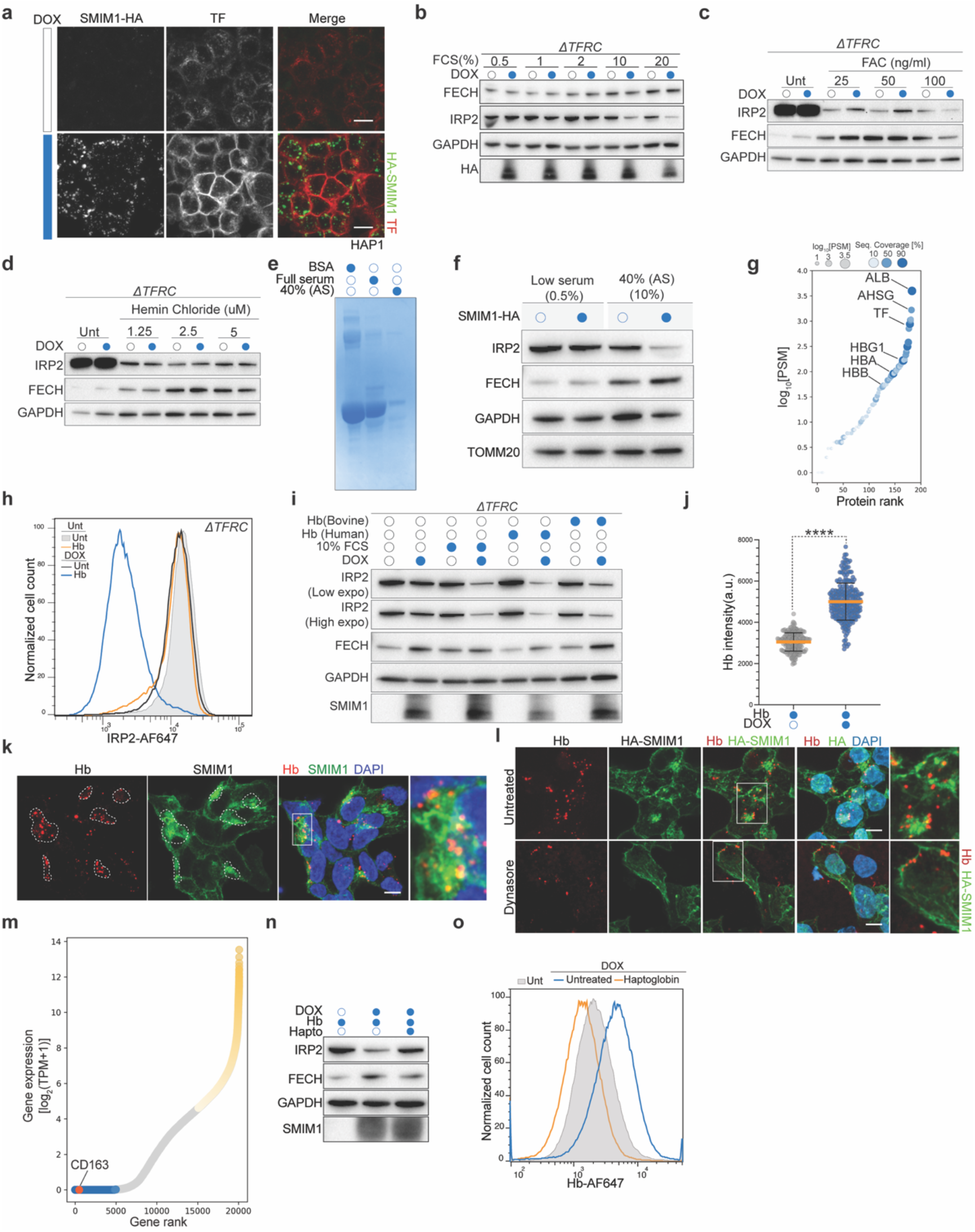
SMIM1 ensures Hb uptake. **(a)** Wild-type HAP1 cells expressing C-terminally HA-tagged SMIM1 following doxycycline induction were live-stained with an anti-HA antibody to detect SMIM1 and labeled transferrin to detect TFRC. The cells were live imaged using a ZEISS LSM 980 Airyscan microscope. Scale bar, 10 μm. (**b**) TFRC knockout HAP1 cells were induced to express SMIM1 with doxycycline (2 µg/mL) for 16 hours, after which the cells were cultured in various concentrations of fetal calf serum (FCS) for an additional 24 hours in the presence of doxycycline. Whole-cell extracts were subjected to immunoblot analysis using the indicated antibodies, with GAPDH serving as a loading control. (**c-d**) TFRC knockout HAP1 cells with doxycycline-inducible SMIM1 were cultured in 0.5% FCS and treated with or without 2 µg/ml doxycycline for 16 hours, followed by either ferric (III) ammonium citrate (FAC) (c) or hemin chloride (d) treatments at the indicated concentrations for an additional 24 hours. Whole-cell extracts were subjected to immunoblot analysis using the stated antibodies, with GAPDH as a loading control. (**e-g**) Mass spectrometry (MS) analysis of the composition of fetal calf serum (FCS) used as a supplement in our media. Full FCS or the supernatant of 40% ammonium sulfate-precipitated FCS was analyzed by denaturing SDS-PAGE and stained with Coomassie (e). The ammonium sulfate supernatant retained the effect on IRP2 levels observed upon SMIM1 expression (f). The composition of 40% ammonium sulfate-precipitated FCS supernatant was analyzed by MS and depicted as a ranking plot (g). (**h**) TFRC knockout cells were treated with doxycycline to induce SMIM1 expression or left untreated as a control for 16 hours in low serum (0.5% FCS), then cultured with 10 µg/ml hemoglobin in low-serum medium for an additional 24 hours in the presence of doxycycline. Cells were subsequently stained for IRP2 and analyzed by flow cytometry. (**i**) TFRC knockout cells were treated with doxycycline for 16 hours to induce SMIM1 expression or left untreated under low-serum conditions, followed by incubation with either 6 µg/ml of bovine or human hemoglobin for another 24 hours under low-serum conditions, with full-serum conditions (10% FCS) included as a control. Whole-cell extracts were analyzed by immunoblotting, with GAPDH as the loading control. **(j)** Hemoglobin intensities were quantified in more than 200 cells per condition for the immunofluorescence staining shown in Fig. 4h. Values represent mean intensity per cell, and *p*-values were calculated using the Mann-Whitney U test. **** p < 0.0001. (**k**) TFRC knockout HAP1 cells were exposed to doxycycline for 24 hours to induce SMIM1 expression, then cultured in the presence of fluorescent hemoglobin (Hb-AF647, 1 µg/ml) for 6 hours. The cells were then fixed, permeabilized, and stained using SMIM1 antibody and DAPI. Scale bar, 5 μm. (**l**) Representative image of the co-localization of fluorescent hemoglobin (Hb-AF647) with SMIM1 in TFRC knockout HAP1 cells untreated or treated with dynasore (20 µM) for 6 hours before fixation. SMIM1 containing a N-terminal HA epitope tag was detected using antibodies directed against HA. Scale bar, 5 μm. (**m**) Gene expression ranking plot in wild-type HAP1 cells using published RNA-sequencing data^37^. Genes were ranked by log_2_(TPM + 1), with TPM representing transcripts per million. The 5,000 highest- and lowest-expressed genes are shown in yellow and blue, respectively, and CD163 is highlighted in red. (**n**) TFRC knockout cells were treated with doxycycline for 16 hours to induce SMIM1 expression or left untreated as a control, followed by incubation with 6 µg/ml human hemoglobin with or without 24 µg/ml haptoglobin for another 24 hours at low serum (0.5% FCS). Whole-cell extracts were subjected to immunoblot analysis using the indicated antibodies, with GAPDH serving as a loading control. (**o**) TFRC knockout cells were exposed to doxycycline for 24 hours to induce SMIM1 expression or left untreated, then cultured in the presence of fluorescent hemoglobin (Hb-AF647, 1 µg/ml) with or without 4 µg/ml of haptoglobin at low serum for an additional 6 hours. The cells were subsequently analyzed by flow cytometry.

**Extended Data Figure 10.**
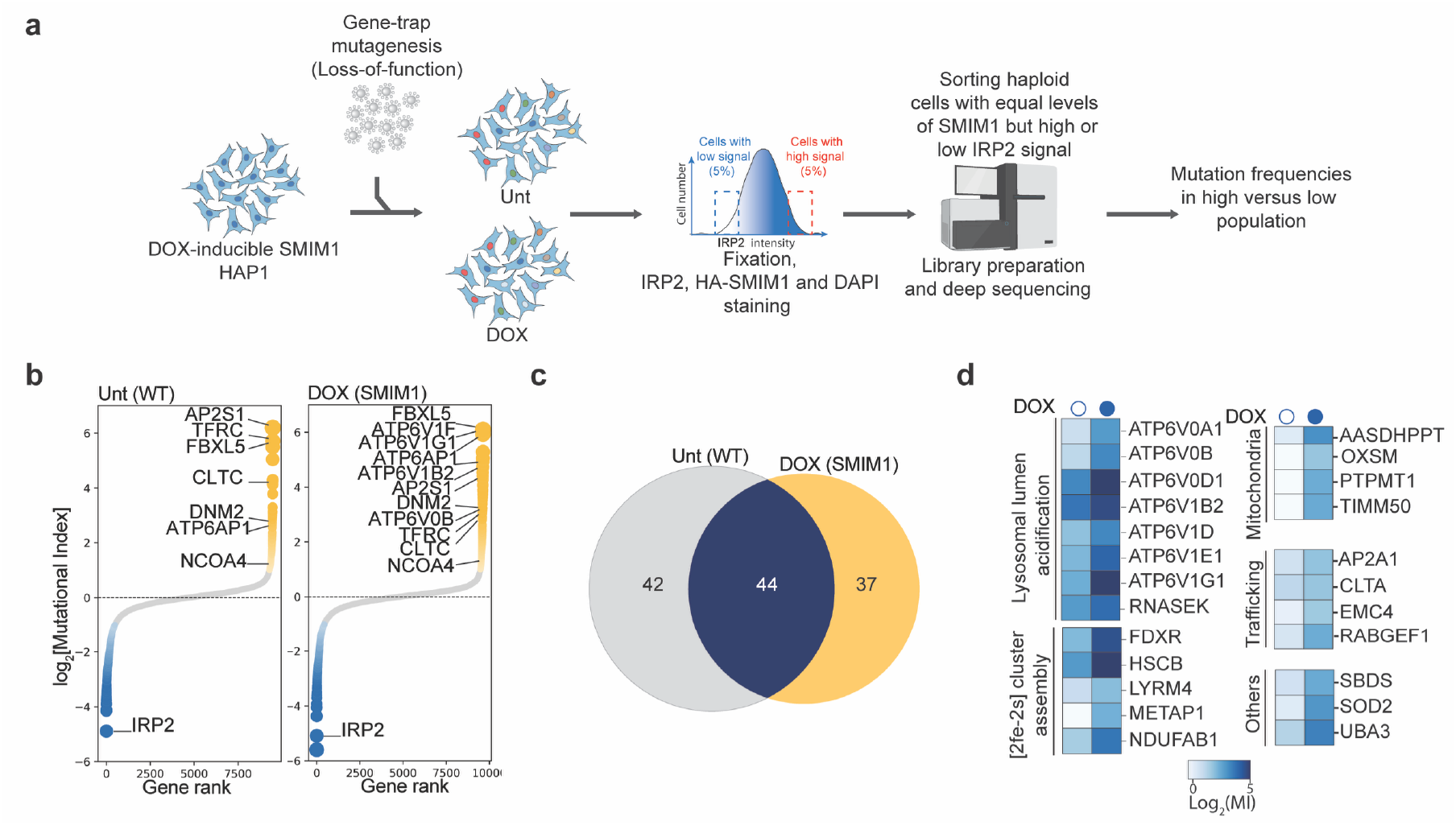
Loss-of-function haploid screens identify SMIM1-specific regulators of Hb uptake. (**a**) Schematic overview of the experimental setup for loss-of-function FACS-based haploid genetic screens using IRP2 as a readout in doxycycline-inducible SMIM1 wild-type HAP1 cells cultured in low serum (0.5% FCS) in the presence of Hb, with or without doxycycline. (**b**) IRP2 screens conducted using loss-of-function haploid genetic screens as described in (a). The data are presented as ranking plots based on mutation index (MI) values. Genes significantly enriched in either channel (two-sided Fisher’s exact test; FDR-corrected p-value ≤ 0.05; total number of unique insertions > 50) are color-coded and top regulators are labeled. (**c**) Venn diagram of factors that strongly increase IRP2 levels (FDR-corrected P ≤ 0.05, log2[MI] < −1, and total unique insertions > 50) identified using loss-of-function genetic screens in SMIM1-expressing versus control cells. (**d**) Heatmap depicting SMIM1-specific regulators organized based on their Gene Ontology terms.

**Extended Data Figure 11.**
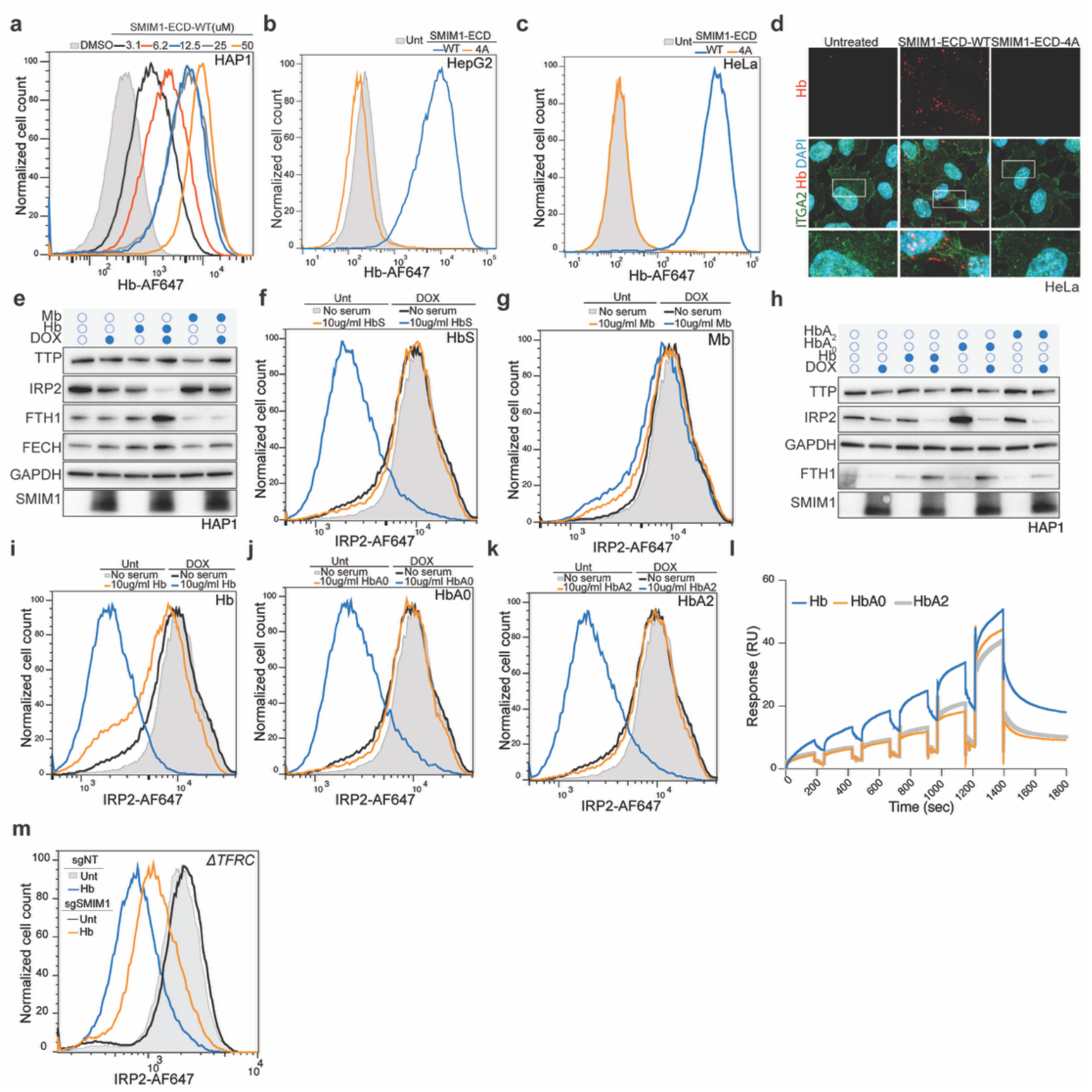
SMIM1 binds different Hb variants and promotes their uptake. (**a**) Wild-type HAP1 cells were incubated either with DMSO or different concentrations of the wild-type extracellular domain of SMIM1 (SMIM1-ECD-WT) for 2 hours with 1 μg/ml labeled Hb (Hb-AF647). The cells were then washed, harvested, and subjected to flow cytometry analysis for Hb fluorescence. (**b-c**) Wild-type HepG2 (b) or HeLa (c) cells were treated for 2 hours with 12.5 μM of either the wild-type extracellular domain of SMIM1 (SMIM1-ECD-WT; C-terminal 15 amino acids of wild-type SMIM1), the HKCK-to-alanine mutant (SMIM1-ECD-4A), or DMSO, in the presence of 1 μg/ml Hb-AF647. Cells were then washed, harvested, and analyzed by flow cytometry to quantify Hb fluorescence. (**d**) HeLa cells were incubated with 12.5 μM of either SMIM1-ECD-WT or SMIM1-ECD-4A, or DMSO for 2 hours in the presence of 1 µg/ml Hb-AF-647. The cells were then subjected to immunofluorescence analysis. (**e**) SMIM1 expression was induced in TFRC knockout HAP1 cells by treatment with doxycycline for 16 hours or left uninduced as a control. Cells were subsequently cultured in low-serum medium (0.5% FCS) supplemented with 10 µg/ml hemoglobin (Hb) or 10 µg/ml myoglobin (Mb) for an additional 24 hours. Whole cell extracts from the conditions were then analyzed by immunoblot using the indicated antibodies, with GAPDH as a loading control. (**f-g**) SMIM1 expression is induced in TFRC knockout HAP1 cells by treatment with doxycycline, or cells were left untreated, for 16 hours. Cells were then cultured in low serum with 10 µg/ml HbS (f) or myoglobin (Mb) (g) for an additional 24 hours. The cells were subsequently stained for IRP2 and analyzed by flow cytometry. (**h**) SMIM1 was induced in TFRC knockout HAP1 cells by doxycycline treatment for 16 hours, or cells were left untreated. Cells were then cultured in low serum for an additional 24 hours with 10 µg/ml of either hemoglobin (Hb), hemoglobin variant A0 (HbA0), or hemoglobin variant A2 (HbA2). Whole cell extracts from the conditions were then analyzed by immunoblot using the indicated antibodies, with GAPDH as a loading control. (**i-k**) Similar conditions as in (h) and cells were stained for IRP2 and analyzed by flow cytometry. (**l**) Surface plasmon resonance (SPR) measurements of binding of human hemoglobin (Hb), hemoglobin variant A0 (HbA0), and hemoglobin variant A2 (HbA2) to the immobilized wild-type extracellular domain of SMIM1 (SMIM1-ECD-WT). (**m**) TFRC knockout HepG2 cells were transduced with a non-targeting sgRNA (sgNT) or sgRNAs targeting SMIM1. After puromycin selection, the cells were cultured with 10 µg/ml hemoglobin in low serum for 48 hours. The cells were subsequently stained using antibody directed against IRP2 or DAPI to stain DNA and analyzed using flow cytometry.

**Extended Data Figure 12.**
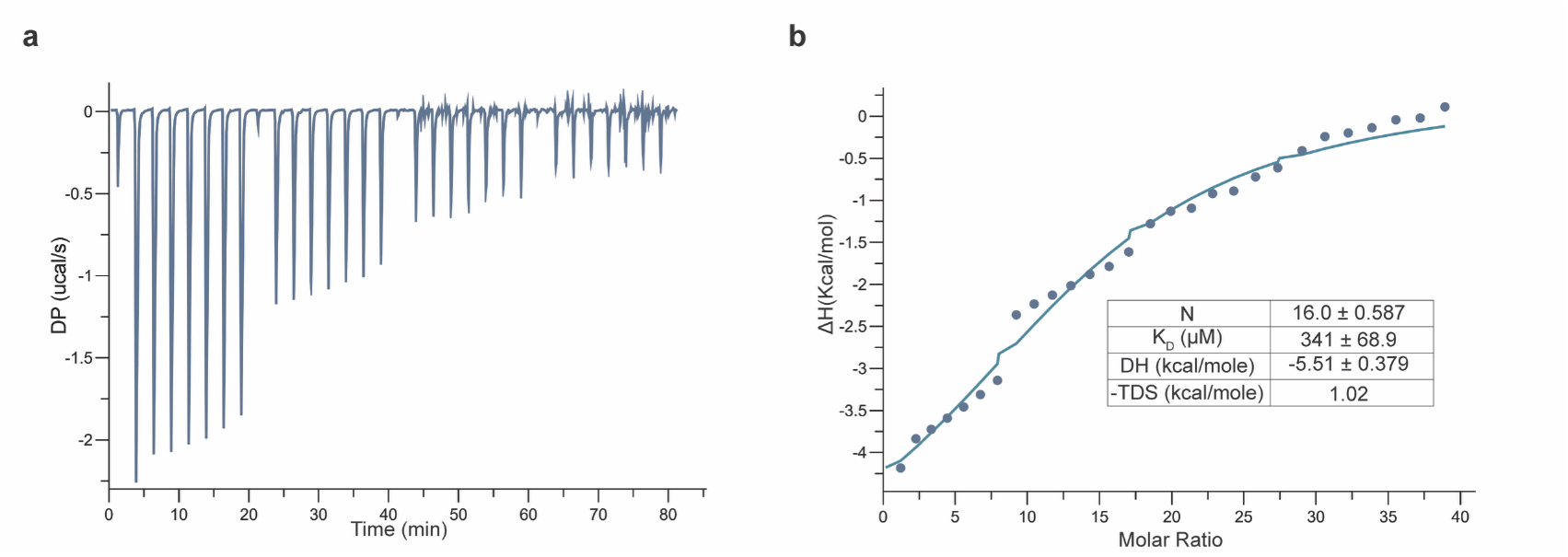
(a-b) SMIM1 binding to Hb using Isothermal Titration Calorimetry (ITC). Binding of SMIM1 to Hb was measured by isothermal titration calorimetry (ITC). Hb (75 µM) was loaded into the cell and SMIM1 (ECD-WT, 3 mM) was titrated into the cell. Control experiments were performed by titrating SMIM1 into PBS.

**Extended Data Figure 13.**
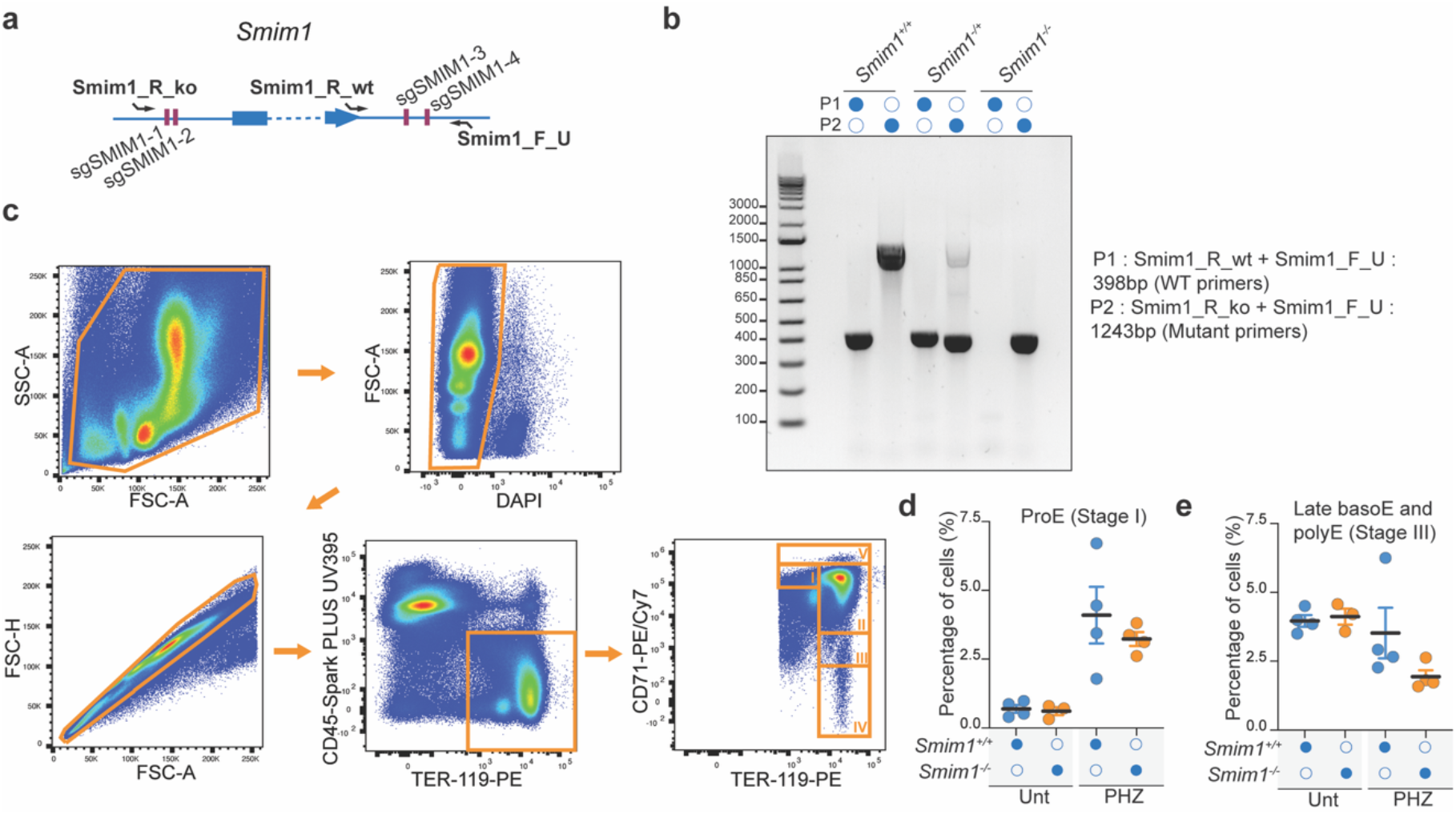
Generation and hematopoietic phenotyping of Smim1 knockout mice. (**a**) sgRNA used to generate Smim1 knockout mice and the strategy for genotyping the mice. (**b**) Genotyping of wild type control (*Smim1*^*+/+*^), *Smim1* heterozygous (*Smim1*^*+/−*^), and *Smim1* homozygous knockout (*Smim1*^*−/−*^) mice. (**c**) Representative gating strategy used to identify bone marrow erythroid populations, including ProE (stage I), BasoE (stage II), late BasoE and PolyE (stage III), OrthoE (stage IV), and CD71^high^ BasoE (stage V). (**d-e**) Frequencies (%) of ProE (d) or late BasoE and PolyE (e) in the bone marrow of *Smim1*^*+/+*^ and *Smim1*^*−/−*^ mice in unchallenged conditions (*n = 4*) or following phenylhydrazine (PHZ)-induced anemia (*n = 4*). Data are shown as mean ± SEM.

**Extended Data Figure 14.**
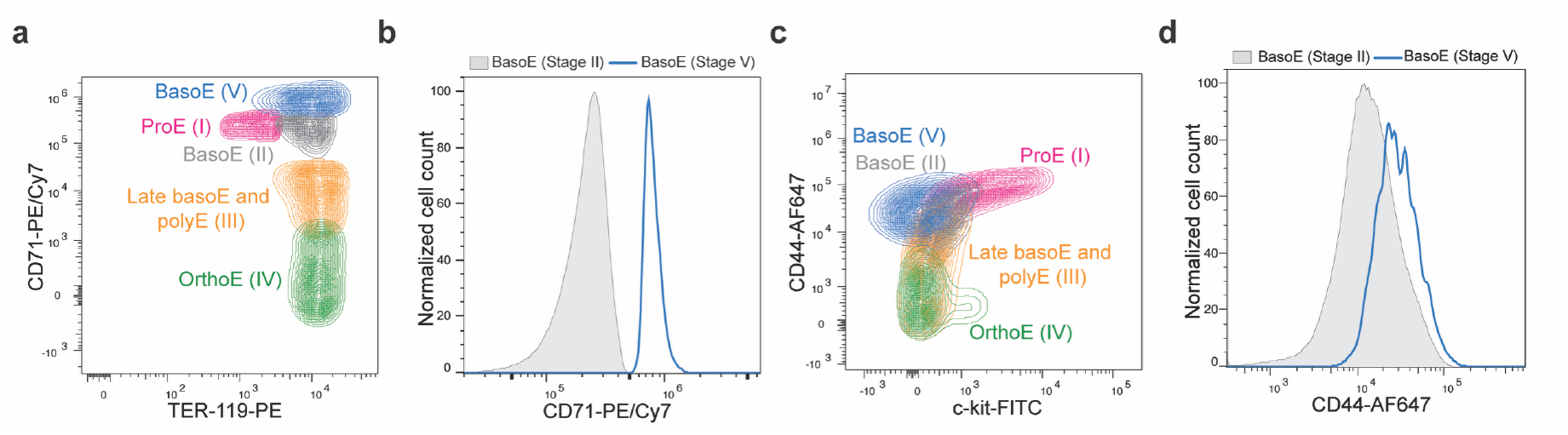
Smim1 knockout mice accumulate basophilic erythroblasts with elevated CD71 expression. (**a**) Flow cytometry density plot of bone marrow erythropoietic populations from a representative Smim1 knockout mouse after PHZ treatment. The plots display CD71 (TFRC) intensity versus TER-119. (**b**) Comparison of BasoE (stage II) and BasoE with high TFRC expression (stage V) based on CD71 surface expression. (**c**) Similar to (a), displaying CD44 versus TER-119. (**d**) Comparison of BasoE (stage II) and BasoE CD71^high^ (stage V) based on CD44 surface expression.

